# A hybrid machine learning framework leveraging biophysicochemical insights for scalable discovery of protein-ligand interactions

**DOI:** 10.1101/2025.03.16.643501

**Authors:** Achal Rayakar, Chaitanya Kumar Jaladanki, Qunxiang Ong, Bin Tan, Junjun Chen, Alicia Qian Ler Tan, Ler Ting Rachel Lim, Navita Kohaal, Yu Chen, Jun Wang, Weiping Han, Jiancheng Hu, Hwee Kuan Lee, Hao Fan

## Abstract

Improving *in silico* compound-protein interaction (CPI) predictability is critical for productive drug discovery. Current deep learning approaches largely rely on end-to-end models trained on limited labeled CPI datasets, overlooking the representational power of large-scale biochemical foundation models. We present COMRADE (Contrastive Multirepresentation Accelerated Docking Engine), a hybrid virtual screening framework that accelerates docking by triaging compounds using CE-Screen (Contrastive Embedding-Screen). CE-Screen leverages seven high-dimensional pretrained representations – including those from protein language models and molecular transformers, along with an original physics-based interaction potential encoding – for rapid first-pass screening ∼100× faster than docking. Its contrastive compression neural network maps these inputs onto a single compact, discriminative representation optimized for CPI prediction via a lightweight ensemble classifier. CE-Screen outperforms state-of-the-art end-to-end models by up to 111.11% on retrospective benchmarks and is successfully used to triage ∼10.8 million compounds against five targets, yielding novel hits for each one – including a new scaffold for the branched-chain ketoacid dehydrogenase kinase (BCKDK), an understudied yet high-value target in metabolic disease and oncology.

---

Compounds that bind to proteins can modulate their activity, either enhancing or inhibiting their function. Identifying such compounds is a critical first step in drug discovery, after which they are assessed for efficacy and safety in humans. The multi-stage preclinical-to-market discovery process is marked by a success rate of ∼1 in 20,000-30,000^1^ and can span over a decade with a cost upwards of USD 1 billion^2–4^. Enhancing the productivity of this process is imperative, especially as multi-billion-compound chemical libraries are now readily accessible for testing against each protein target of interest^5^. Compounding the challenge, the druggable human genome is estimated to contain approximately 4,479 protein-coding genes^6^, each representing a potential therapeutic target. A key avenue for improving this productivity lies in the ability to identify more ligands at a faster pace.

Although routinely used for ligand discovery, high-throughput screening methods that rely on biochemical assays are time- and cost-inefficient for directly screening large libraries. These methods are limited to testing only thousands of compounds per week^7^ while consuming substantial amounts of reagents. To overcome this bottleneck, force-field-based molecular docking has conventionally been a key *in silico* approach for prioritizing promising compounds from chemical libraries for experimental validation. However, even advanced docking techniques are too slow for exhaustive screening of modern ultra-large-scale chemical libraries; conducting such screenings within practical timeframes requires extensive parallel computing resources, which can cost hundreds of thousands of USD^8^.

Recent advances in machine learning (ML), particularly within deep learning, have significantly enhanced virtual screening^9^. For instance, graph convolutional networks (GCNs) excel in capturing complex molecular topologies and electronic interactions without requiring detailed force-field calculations by processing molecules as graphs, where atoms are represented as nodes and bonds, edges. This allows GCNs to efficiently learn chemically-rich representations of compounds, enabling rapid screening while bypassing the computational cost of traditional docking approaches^10^. These advancements deliver both high performance and speed, making virtual screening practical for ultra-large chemical libraries.

In parallel with the development of deep learning methodologies for compound-protein interaction (CPI) prediction, sophisticated representations of both protein targets and small molecules have emerged that leverage large-scale datasets and innovative model architectures. Among the latter, transformer-based models have risen to prominence. By employing self-attention mechanisms, transformers excel at learning complex contextual relationships within input data, enabling a deeper understanding of sequential information^11^. For protein sequences, transformers can excel at, for example, capturing long-range dependencies between amino acids, enabling the extraction of highly informative embeddings that accurately reflect structural, functional, and evolutionary properties^12^. For small molecules, transformers trained on SMILES strings can embed chemical structures into highly organized latent spaces that cluster similar molecules together, capturing subtle relationships between atomic arrangements and chemical properties^13^.

Despite these advances, prevailing state-of-the-art CPI prediction models completely or in large part employ end-to-end deep learning^14–19^. These approaches internally derive compound-protein representations directly from low-level inputs (e.g., protein sequences and SMILES) and use them to predict interactions. While performant, end-to-end models rely on CPI datasets that are orders of magnitude smaller than the large, domain-specific unlabeled datasets used to train specialized foundation models. As a result, end-to-end models may be unable to capture the rich, multi-range biochemical relationships large-scale pretraining can expose. By not incorporating these pretrained representations, therefore, these models may be less effective in virtual screening tasks, particularly for targets with limited or no known interaction data.

Here, we present COMRADE (Contrastive Multirepresentation Accelerated Docking Engine), a virtual screening framework that expedites docking with the rapid, accurate triage of large chemical libraries using our ML CPI classifier, CE-Screen (Contrastive Embedding-Screen). CE-Screen leverages several informative feature sets created by specialized methods, including language model embeddings and an original interaction potential encoding that serves as a physics-based advancement over existing efforts. CE-Screen also circumvents the significant pitfalls associated with modelling CPIs using high-dimensional and heterogeneous feature sets through its use of a feature compression neural network. This network employs a modified contrastive loss function to learn to consolidate its input feature sets into a unified, compact, and discriminative compound-protein representation optimized for CPI prediction with CE-Screen’s classification head, an ensemble of randomized decision trees. With this hybrid, two-module architecture, CE-Screen outperforms end-to-end alternatives by up to 111.11% and 23.39% (per AUPRC and logAUC, respectively) on two retrospective benchmarking datasets constructed for this work. These datasets, designed to assess models as universal CPI classifiers, feature a broad range of protein families and diverse compounds. Moreover, despite 98% of their compounds being non-interacting, CE-Screen consistently uncovers over five ligands among its top 10 predictions – while being ∼100× faster than standard-precision molecular docking. Finally, COMRADE is used to screen a ∼10.8 million-compound library against five targets, including two absent in its training data; it successfully identifies one novel scaffold for each target, including the branched-chain ketoacid dehydrogenase kinase, an understudied yet high-value target in metabolic disease and oncology. We therefore propose COMRADE as a modular, scalable, and rapid representation-driven ligand discovery framework.

## Results

### The COMRADE framework

COMRADE is a four-stage virtual screening framework designed to rapidly predict CPIs across large chemical libraries while minimizing the need for molecular docking. The first three stages, collectively referred to as CE-Screen, integrate contrastive neural network-based feature compression with decision tree-based CPI classification. To predict CPIs, CE-Screen processes 2,457 features representing the compound-protein pair of interest. This high-dimensional input is drawn from seven diverse methods to provide a comprehensive characterization of the protein target and compound (Fig. 1a).

**Figure 1:**
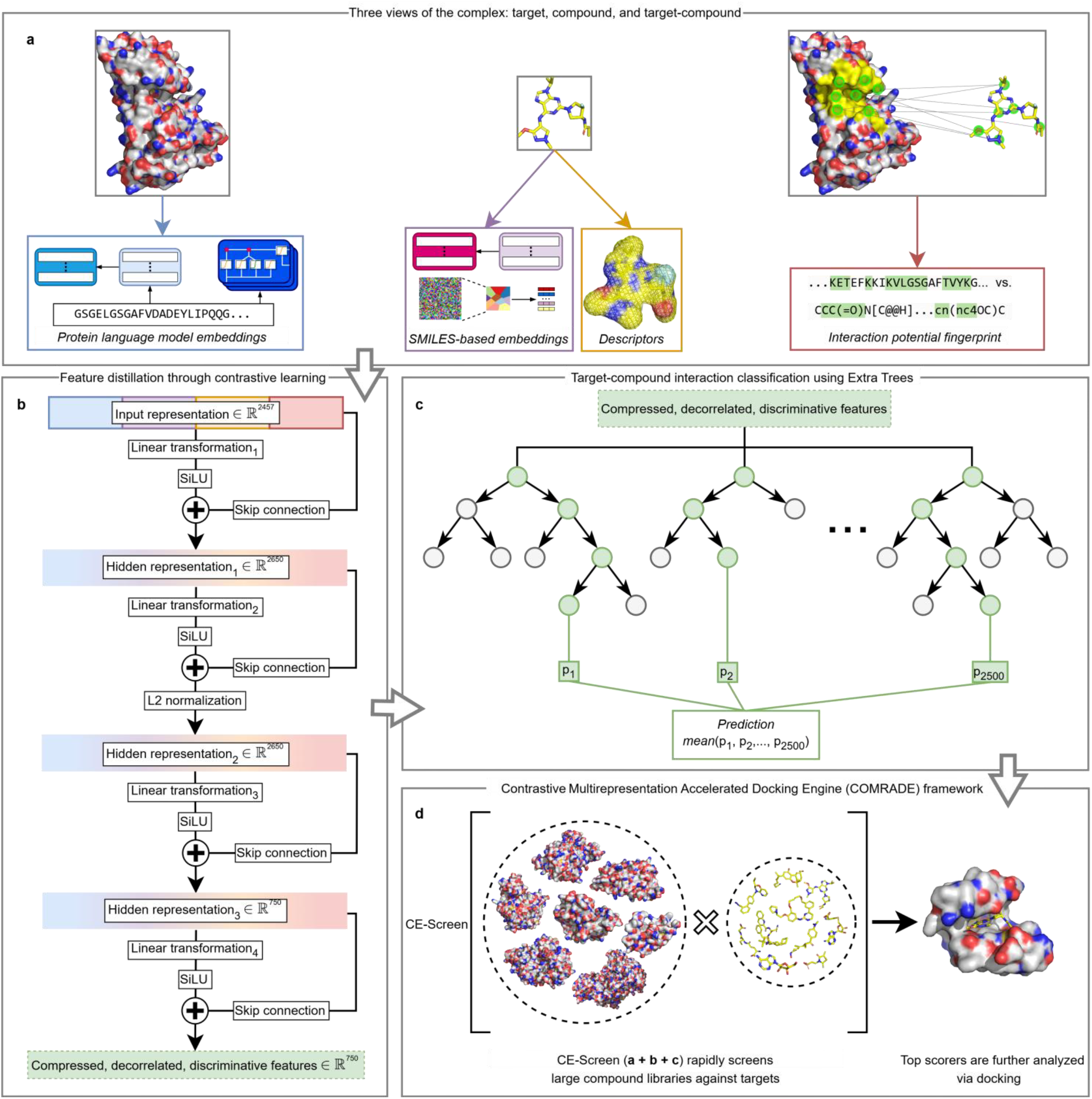
The COMRADE framework -- CE-Screen followed by docking. **a**, CE-Screen represents compound-protein pairs along three feature categories: protein sequence embeddings; compound embeddings and descriptors; and an interaction potential encoding based on Merck Molecular Force Field atom types present in the target pocket and compound. **b**, the resulting 2,457-dimensional feature vector is compressed into a 750-dimensional representation using supervised contrastive learning with a residual neural network. This network is trained to separate interacting (ligand) and non-interacting (decoy) pairs while promoting feature decorrelation. **c**, the discriminative, compressed representation is passed to an Extra Trees ensemble classifier, which outputs a pseudoprobability of interaction. **d**, CE-Screen (**a**-**c**) is used to rapidly score large chemical libraries to identify the most promising compounds for further investigation via more computationally-intensive molecular docking.

Two of the constituent feature sets are embeddings derived from protein language models (pLMs) which convert protein sequences into per-amino acid distributed vector representations^20^. Specifically, CE-Screen uses outputs from DLM-LSTM and ESM-2^21^. Both models employ a self-supervised masked language modeling approach^22^ to produce representations that reflect proteins’ structural and functional characteristics and capture both local and global sequence dependencies. Trained on large-scale unlabeled data, such as the 48 million protein sequences used for ESM-2, these models encode a broad contextual foundation within their representations. Prior to use by CE-Screen, however, each model’s per-amino acid representations are aggregated into a single fixed-dimensional vector via a weighted mean, with greater weight assigned to representations corresponding to residues within the target pocket according to a capped inverse power function (see Methods for further details).

Four other feature sets encode compounds based on their SMILES strings^23^: those derived from MolFormer-XL^24^, Mordred^25^, mol2vec^26^, and MACAW^27^. MolFormer-XL is a large-scale language model trained using masked language modeling on hundreds of millions of SMILES strings to produce molecular embeddings that capture both local substructural details and global molecular context. Mordred generates traditional molecular descriptors, capturing features such as atom counts, bond connectivity, adjacency matrices, aromaticity, and topological indices. mol2vec improves upon traditional molecular fingerprints through aggregating compounds’ substructural embeddings, where these embeddings are learned by the method to reflect properties such as substructures’ presence, type, relative importance, co-occurrence patterns, and chemical similarity. Finally, MACAW embeds compounds into a low-dimensional, continuous space, arranged by the method using reference compounds so as to locate structurally and physicochemically similar compounds closer together.

The last feature set, and one introduced in this work, is an interaction potential encoding (IPE) that captures target-compound compatibility using physics-based atom typing. Prominent approaches, such as that employed by RF-Score-VS^28^, encode intermolecular interactions by counting atom-atom contacts in resolved or docked target-compound structures. However, since resolved structures for compounds of interest are often unavailable, and docking is computationally expensive, the IPE circumvents these requirements by relying solely on the Merck Molecular Force Field (MMFF) atom types^29^ of the target and compound. Unlike conventional encodings that rely primarily on elemental identity, MMFF types classify atoms based on their hybridization state, bonding environment, and chemical context.

IPEs are generated in 4 steps. First, an MMFF type is assigned to each atom in the target’s binding pocket and in the compound. Next, the MMFF types of the atoms covalently bonded to these atoms are identified. This step enables the construction of an MMFF type pair for each atom in the binding pocket and the compound, where the first element of the pair represents the atom’s own type and the second represents the types of its neighboring atoms. In the third step, the frequencies of unique combinations of target and compound MMFF type pairs are computed. Lastly, principal component analysis is applied to consolidate the large number of pair counts into a concise feature set. By integrating both immediate and broader atomic environments, the resulting encoding provides a comprehensive representation of the interaction potential between the target and compound.

The seven feature sets are concatenated into a 2,457-dimensional vector representing the compound-protein pair. While informative, this representation is high-dimensional relative to the size of CPI training datasets, which typically contain only a few thousand to tens of thousands of labeled pairs. A purely classical ML approach that trains CE-Screen’s Extra Trees^30^ classification head – which works by partitioning the feature space into regions with similar outcomes – on this representation directly would be at risk of overfitting due to the sparsity of training pairs within this high-dimensional feature space^31^. Additionally, the accumulation of noise across many features in high-dimensional settings can obscure discriminative data patterns, further hindering the classifier’s ability to predict the interaction of compound-protein pairs not featured in the training data^32^. A contemporary approach of using a deep neural network for classification would also face these challenges. In particular, its end-to-end training would force the network’s entire capacity to directly optimize classification accuracy, potentially driving its encoder to discard input signals that do not immediately reduce classification loss. Such specialization can lead the model to overfit to dataset-specific or label-specific patterns, particularly in data-limited and variegated settings like CPI prediction.

By contrast, CE-Screen mitigates these risks by adopting a compression-and-classification architecture that stages learning into two distinct steps. First, its contrastive neural network compresses and structures the feature space by learning to broadly separate interacting and non-interacting pairs, without committing to the specific decision boundaries of any one classifier. This compression is still guided by the same interaction labels as classification is, but the contrastive objective focuses on global class separation in the network’s output embedding space rather than directly minimizing classification error. This broader objective helps ensure that the resulting compressed representations retain a generalizable, class-discriminative structure that is less tied to dataset-specific idiosyncrasies. CE-Screen then defers classification to an Extra Trees ensemble trained on these contrastive embeddings, reducing its risk of overfitting while benefiting from the ensemble’s robustness to small samples.

Specifically, CE-Screen uses a residual neural network^33^ to map its 2,457-dimensional input onto a compact 750-dimensional embedding space (Fig. 1b). Skip connections are included in the network to enable incremental adjustments to its learned internal representations, discouraging the network from making unnecessary or overly complex transformations. This network is trained using a modified supervised contrastive loss^34^ introduced in this work, which incorporates a correlation penalty to promote embedding feature diversity. Specifically, the correlation penalty encourages the network to produce features that are orthogonal, which is intended to enhance the performance of the downstream Extra Trees classifier by increasing the diversity of its constituent decision trees (Fig. 1c).

Extra Trees, as an ensemble of independently trained decision trees, benefits from decorrelated input features because each tree is built using a random subset of features. When features are highly correlated, different trees may still rely on similar information, leading to overlapping decision boundaries and reduced ensemble diversity. Decision trees classify inputs by making hierarchical, axis-aligned splits; if many trees use correlated features, they tend to construct similar splits, which limits the ensemble’s ability to capture distinct patterns. This redundancy also weakens the variance-reduction advantage typically provided by ensembling^35^. By promoting feature decorrelation, the contrastive compression network encourages structural diversity across trees, yielding more complementary decision boundaries that can enhance CE-Screen’s generalization. Furthermore, CE-Screen’s large ensemble size – featuring 2,500 trees – further stabilizes predictions by ensuring that no single tree overly influences the final output.

Namely, CE-Screen’s Extra Trees classifier creates an interaction prediction for each input compound-protein pair based on a majority vote among its constituent decision trees. Each tree independently classifies the pair as either interacting (output: 1) or non-interacting (0). While this enables a consensus binary classification by the ensemble, the mean of these votes also provides a confidence score (or pseudoprobability) of interaction. These pseudoprobabilities are used within the COMRADE framework to prioritize compounds for further evaluation, with only those having the highest scores being selected for downstream investigation and interpretation via docking (Fig. 1d). LigPrep^36^ is used to preprocess these compounds, generating all possible ionization states at pH 7.0 ± 2.0 and multiple conformers per compound. Targets’ structures are prepared by assigning hydrogen atoms and partial charges, and receptor grids are generated around bound ligands using Glide’s Receptor Grid Generation tool. Docking grids are defined to encompass entire active sites with sufficient margins to ensure the inclusion of all relevant binding interactions. Prepared compounds are then docked to their target using the Glide in its Standard Precision mode (Glide-SP)^37–39^. After docking, top-ranked compounds (as determined by their docking scores) have their docked poses visually inspected, and the most promising candidates are experimentally tested.

### Distinct, diverse retrospective testing datasets

Three datasets are used to train and retrospectively test CE-Screen against other CPI classification models: a deduplicated version of the Directory of Useful Decoys, Enhanced^40^ (D-DUD-E), and original subsets of the BindingDB^41^ and ChEMBL^42^ databases (Table 1). Testing follows a cascading approach: first, models are trained on D-DUD-E and tested on BindingDB; next, they are trained on both D-DUD-E and BindingDB and tested on ChEMBL. The BindingDB and ChEMBL subsets introduced in this work are designed to be significantly distinct from each other and from D-DUD-E, ensuring that these tests provide a dual, robust assessment of the models’ abilities to predict the interaction of diverse compound-protein pairs. Specifically, both datasets have been created to present two challenges: identifying whether novel compounds interact with targets similar to those encountered in model training (i.e., whether they are ligands instead of decoys), and identifying novel ligands for novel targets. The first scenario tests their ability to learn underlying interaction principles rather than overfitting to spurious correlations from target-ligand pairings in training. The second measures models’ generalization ability across both protein and chemical diversity when there is little precedent in the training data to rely on. Together, these challenges reward only models capable of both fine-grained interpolation and extrapolation from their training data.

**Table 1:** datasets used for model training and retrospective testing. On average, D-DUD-E, BindingDB, and ChEMBL have 201, 154, and 51 ligands per target (respectively). The datasets also feature decoys to serve as complementary examples of non-binding compounds. In testing, BindingDB and ChEMBL include 50 times the number of decoys as there are ligands to reflect ligands’ practical rarity. In training, D-DUD-E and BindingDB maintain parity between ligands and decoys to aid model learning. While D-DUD-E and ChEMBL are only ever used for model training and testing respectively, BindingDB plays both roles. Its training form is derived by resampling one-fiftieth of the decoys within its testing configuration while retaining the same targets and ligands.

| Dataset | Targets | Ligands | Decoys |  |
| --- | --- | --- | --- | --- |
|  |  |  | In training | In testing |
| Deduplicated DUD-E (D-DUD-E) | 102 | 20,456 | 20,456 | NA; only used in training. |
| BindingDB subset (BindingDB) | 49 | 7,565 | 7,565 | 378,250 |
| ChEMBL subset (ChEMBL) | 22 | 1,114 | NA; only used in testing. | 55,700 |

Given the scarcity of experimentally-verified weakly-binding and non-binding compounds to the targets of interest, algorithmically-selected decoys must be used as stand-ins during model training and testing. For each dataset, two types of decoys are compiled: matched decoys and randomly-assigned decoys. The matched decoy selection style, as introduced in the creation of DUD-E, selects compounds from the ZINC database^43^ that are physicochemically similar but structurally distinct from their corresponding ligands, with a T_c_ typically between 0.20 and 0.40. This process is designed to create challenging decoys that are unlikely to be false negatives, enabling models to be trained and tested to discern the structural nuances underpinning ligand binding. However, by design, matched decoys may fail to capture the extensive diversity of screening libraries that trained models are practically intended to be applied to^44^. This work therefore introduces and also employs a second, random decoy selection method. In this method, compounds from ZINC are randomly assigned as decoys to each ligand of every target. Namely, ZINC organizes its compounds into tranches based on logP and molecular weight, and since some tranches are larger than others, simple random sampling would inadequately represent compounds from less common tranches. To address this, the random sampling procedure is stratified by tranche to promote physicochemical diversity among decoys. To also promote structural diversity, decoys are selected to have a T_c_ of at most 0.30 with any other decoy for the same target. Finally, since decoys are selected independently for each target, any duplicates across targets are replaced with new, randomly chosen compounds. Both matched and random decoys offer a limited and biased approximation of the ideal weak-/non-binder compound set associated with each target, which would be large, diverse, and experimentally-verified. Nonetheless, the limitations associated with both decoy types are distinct: training and retrospectively testing with them, therefore, provides complementary perspectives of models’ utility. With random decoys, models learn and are evaluated on their ability to identify ligands amidst a broad, heterogeneous chemical space. While this is a coarser-grained challenge than that posed by matched decoys, considering both sets of tests in unison can help determine which models have both a superior capacity to generalize across diverse chemical environments and to also learn the narrow distinctions inherent in the matched decoy design.

Moreover, existing CPI classification studies typically employ retrospective testing datasets with 1-10 decoys per ligand^14–19,45–47^. While the true ratio of weak- and non-binders to ligands in chemical libraries is both practically difficult to ascertain and varies by library and target, these conventional ratios may fail to capture the scarcity of ligands in real-world screening – where they often represent as little as 0.01% of the library^48^. Accordingly, following the approach used in the creation of DUD-E, a decoy-to-ligand ratio of 50 is adopted for each dataset to better approximate this scarcity, although even this ratio may be optimistic in certain cases.

Given the limitations inherent to retrospective testing, the prospective performance of the COMRADE framework is also evaluated on five targets: three human kinases from the D-DUD-E dataset – the serine/threonine-protein kinase B-Raf (BRAF), epidermal growth factor receptor (EGFR), and Mitogen-activated protein kinase (MEK1) – along with the SARS-CoV-2 main protease (M^pro^) from ChEMBL, and the branched-chain ketoacid dehydrogenase kinase (BCKDK), a target absent from the three datasets with only seven known ligands^49–52^. This is done by first using CE-Screen, trained on D-DUD-E and BindingDB with random decoys, to score (i.e., generate pseudoprobabilities for) ∼10.8 million ZINC compounds against each target. The 10,000 top-scoring compounds for each target are docked and inspected, with a manually-selected ∼10-20 then being experimentally validated. Random decoys are selected over matched decoys for CE-Screen’s training to enable it to better distinguish between the unknown ligands to be identified and the large, diverse background of weak-/non-binders among the compounds being scored. This experimental setup therefore serves as an evaluation of COMRADE’s ability to identify promising hits for both well- and under-characterized targets.

### CE-Screen achieves superior results in retrospective testing

CE-Screen is benchmarked against five ML models and Glide-SP (used as a control) on BindingDB and ChEMBL, using both matched and random decoys. These ML models represent the current state-of-the-art in CPI classification and are selected to be architecturally diverse, employing a variety of convolutional neural network^53^, graph neural network^54^, and transformer^11^ (including pLM) modules. Notably, therefore, none of these representative top performers employ classical machine learning. Moreover, these models learn protein and compound representations in an end-to-end fashion, completely or in large part forgoing manual feature engineering.

Each model’s performance is primarily compared using the logAUC metric, which is the area under the curve measuring ligand recall as a function of the logarithm of the dataset coverage^55^. Here, dataset coverage refers to the proportion of the dataset’s top-ranked compound-protein pairs considered, sorted in descending order by their pseudoprobabilities of interaction as predicted by the model; this axis is transformed in calculating this metric to emphasize models’ early enrichment performance. Table 2 displays that CE-Screen consistently outperforms its alternatives across all testing dataset-decoy style scenarios. With random decoys, CE-Screen achieves a logAUC improvement of 8.65% over the next-best model on BindingDB and 6.55% on ChEMBL. This lead widens with matched decoys, increasing to 23.39% and 8.92% on BindingDB and ChEMBL respectively. Additionally, CE-Screen outperforms Glide-SP by over 100% while being approximately 100 times faster, as Glide-SP requires about ten seconds to score each compound-protein pair^56^. By narrowing docking efforts to high-confidence candidates identified by CE-Screen, the COMRADE framework therefore offers a reliable, scalable, and cost-effective solution to accelerating ligand discovery workflows.

**Table 2:** model performance under each testing dataset and decoy style scenario (best, <u>second best</u>). To assess variability in ML model training due to the random initialization of parameters and order of training data, testing was conducted four times with different random seeds. The mean and standard deviation of these runs’ performances are calculated and presented below. In this CPI classification scenario with 50 decoys per ligand, a random classifier yields a logAUC of ∼14.50 (equivalent to an AUROC of 0.50 and AUPRC of ∼0.02), while a perfect classifier achieves a logAUC of ∼71.30 (equivalent to an AUROC and AUPRC of 1.00).

| Model | Matched decoys |  |  | Random decoys |  |  |
| --- | --- | --- | --- | --- | --- | --- |
|  | LogAUC | AUROC | AUPRC | LogAUC | AUROC | AUPRC |
| **BindingDB testing** |  |  |  |  |  |  |
| CE-Screen | **$49.69 \pm 0.33$** | **$0.91 \pm 0.00$** | **$0.38 \pm 0.01$** | **$61.02 \pm 0.14$** | **$0.96 \pm 0.00$** | **$0.65 \pm 0.01$** |
| BridgeDPI<sup>15</sup> | <u>$40.27 \pm 1.35$</u> | <u>$0.86 \pm 0.01$</u> | <u>$0.18 \pm 0.01$</u> | <u>$56.16 \pm 0.31$</u> | <u>$0.93 \pm 0.00$</u> | <u>$0.52 \pm 0.01$</u> |
| DrugBAN<sup>17</sup> | $37.76 \pm 2.12$ | $0.83 \pm 0.02$ | $0.17 \pm 0.02$ | $52.92 \pm 1.51$ | $0.92 \pm 0.01$ | $0.44 \pm 0.01$ |
| PSICHIC<sup>19</sup> | $37.26 \pm 0.85$ | $0.83 \pm 0.01$ | $0.16 \pm 0.02$ | $54.62 \pm 0.79$ | $0.91 \pm 0.01$ | $0.50 \pm 0.01$ |
| DLM-DTI<sup>57</sup> | $33.26 \pm 1.49$ | $0.78 \pm 0.02$ | $0.15 \pm 0.02$ | $43.50 \pm 0.68$ | $0.86 \pm 0.01$ | $0.26 \pm 0.01$ |
| MolTrans<sup>14</sup> | $32.24 \pm 0.83$ | $0.78 \pm 0.01$ | $0.09 \pm 0.01$ | $44.22 \pm 0.97$ | $0.86 \pm 0.01$ | $0.25 \pm 0.01$ |
| Glide-SP | 23.50 | 0.61 | 0.11 | 26.47 | 0.64 | 0.12 |
| **ChEMBL testing** |  |  |  |  |  |  |
| CE-Screen | **$55.44 \pm 0.26$** | **$0.93 \pm 0.00$** | **$0.52 \pm 0.01$** | **$56.11 \pm 0.11$** | **$0.93 \pm 0.00$** | **$0.54 \pm 0.01$** |
| BridgeDPI | $49.51 \pm 0.99$ | <u>$0.90 \pm 0.01$</u> | $0.36 \pm 0.02$ | $52.06 \pm 0.23$ | <u>$0.92 \pm 0.01$</u> | $0.42 \pm 0.01$ |
| DrugBAN | $45.67 \pm 1.81$ | $0.88 \pm 0.02$ | $0.28 \pm 0.03$ | $49.60 \pm 2.52$ | $0.90 \pm 0.02$ | $0.36 \pm 0.01$ |
| PSICHIC | <u>$50.90 \pm 0.89$</u> | <u>$0.90 \pm 0.01$</u> | <u>$0.39 \pm 0.03$</u> | <u>$52.66 \pm 1.01$</u> | $0.90 \pm 0.01$ | <u>$0.45 \pm 0.01$</u> |
| DLM-DTI | $36.42 \pm 1.11$ | $0.80 \pm 0.01$ | $0.16 \pm 0.02$ | $40.42 \pm 1.18$ | $0.85 \pm 0.01$ | $0.20 \pm 0.01$ |
| MolTrans | $34.26 \pm 1.39$ | $0.79 \pm 0.02$ | $0.12 \pm 0.02$ | $41.70 \pm 0.36$ | $0.84 \pm 0.01$ | $0.21 \pm 0.01$ |
| Glide-SP | 25.76 | 0.66 | 0.10 | 25.72 | 0.64 | 0.10 |

Only CE-Screen and Glide-SP show themselves able to perform approximately as well with matched decoys as they do with random decoys. On ChEMBL testing, CE-Screen and Glide-SP exhibit minimal logAUC reductions of 1.21% and -0.15% (respectively) with matched as opposed to random decoys, while the five competing ML models experience larger reductions of 3.46-21.72%. On BindingDB testing, however, these models show logAUC reductions of 37.16-46.59% from random to matched decoys; in contrast, CE-Screen’s and Glide-SP’s logAUC values decrease far less – by (only) 22.80% and 11.22% respectively. Relative to ChEMBL testing, BindingDB testing appears to exacerbate the challenge matched decoys pose over random ones. CE-Screen’s substantial lead over all other models on BindingDB testing with matched decoys is therefore a demonstration of its greater generalizability. The particular contribution of CE-Screen’s distinctive two-module compression-and-classification architecture to the model’s lead over alternative approaches and specifically, robustness against overfitting will be further considered in Table 3.

**Table 3:** experimentally-validated hits identified by COMRADE.

| Target | Hit ZINC ID | Binding affinity (IC<sub>50</sub>, μM) |
| --- | --- | --- |
| MEK1 | ZINC0143157631 | 30.70 |
| EGFR | ZINC1772617172 | 124.60 |
| BRAF | ZINC0489458680 | 37.90 |
| M<sup>pro</sup> | ZINC0100368890 | 31.34 |
|  | ZINC0001224796 | 34.70 |
|  | ZINC0001326077 | 50.26 |
| BCKDK | ZINC7001415662 | 4.10 |

While the rank order of models is nearly identical in terms of the more commonly-used AUROC (area under the recall-false positive rate curve), the metric tends to minimize performance differences between models. On BindingDB testing, CE-Screen’s AUROC improvement over the next-best model is 3.23% with random decoys and 5.81% with matched decoys. On ChEMBL testing, these figures further reduce to 1.09% with random decoys and 3.33% with matched decoys. CE-Screen’s AUROC improvements are far less pronounced than those observed in terms of logAUC as a consequence of AUROC’s replacement of database coverage with the false positive rate. Moreover, this results in an inflated perception of each model’s performance in this imbalanced CPI classification context; for example, on ChEMBL with random decoys, CE-Screen and BridgeDPI both achieve near-perfect AUROC scores of 0.93 and 0.92, respectively. However, their logAUC scores, 56.11 for CE-Screen and 52.06 for BridgeDPI (out of ∼71.30), present both a more moderate and differentiated picture of their performance. Given non-binders significantly outnumber ligands both in retrospective testing and prospective ligand discovery, practically useful models need to achieve both high ligand recall and precision among top-scoring compound-protein pairs.

While the AUPRC (area under the precision-recall curve) metric, also tabulated, accounts for precision, it does not over-weight models’ early enrichment but rather quantifies their enrichment across the entire ranked dataset equally. Consequently, the metric exaggerates performance differences between models relative to logAUC: with random decoys, CE-Screen achieves an AUPRC improvement of 25.00% over the next-best model on BindingDB and 20.00% on ChEMBL. This lead widens dramatically with matched decoys, increasing to 111.11% on BindingDB and 33.33% on ChEMBL. However, in prospective virtual screening applications, where only a small subset of top-ranked ligand candidates can practically be experimentally tested, AUPRC differences overstate those in models’ practical utility.

Along with its logAUC, CE-Screen’s true positives_10_, which counts the number of ligands among the ten highest-scoring (highest-pseudoprobability) compounds for each target, further evidences its high early enrichment ability: the mean true positives_10_ associated with each protein family is usually five or greater (Fig. 2a). In particular, CE-Screen achieves high enrichment even on protein families underrepresented in its training data; Fig. 2b shows that its logAUC values on protein families constituting less than 2% of the training data (e.g., lyases and transporters) are comparable and sometimes even greater than those on kinases which represent about a quarter of CE-Screen’s training. Moreover, variation in familywise logAUC usually occurs only within a 10-point range.

**Figure 2:**
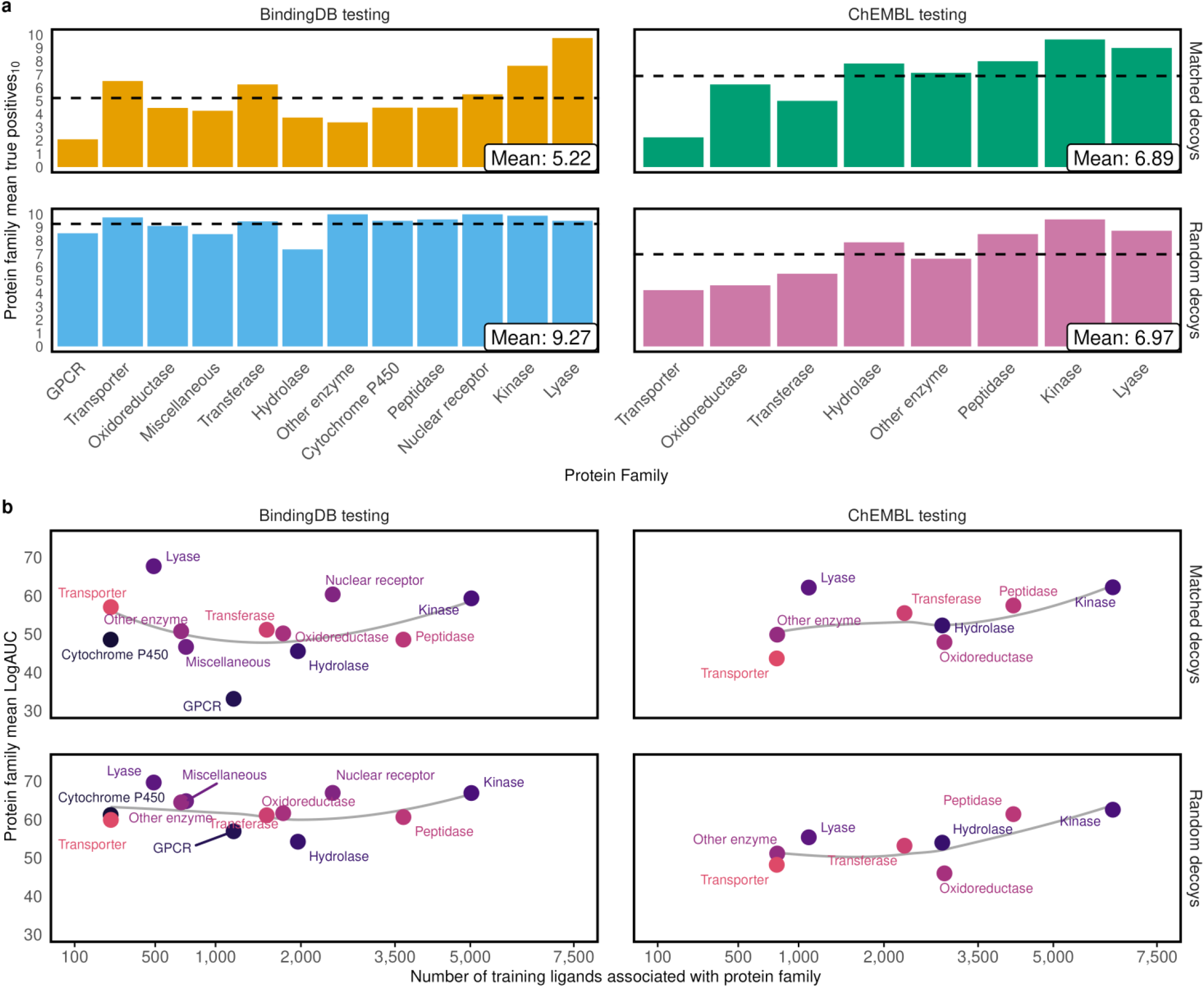
CE-Screen’s performance on each protein family in BindingDB and ChEMBL. **a**, the number of ligands that CE-Screen recovers among the top ten highest-scoring compounds (true positives_10_) under every testing dataset-decoy style scenario. True positives_10_ values are calculated for each target individually before their mean is taken and displayed by protein family. **b**, CE-Screen’s mean logAUC on each protein family plotted against the family’s representation in the training data, for each scenario. While CE-Screen achieves some of its highest logAUC values on the best-represented protein families (peptidases and kinases), the relationship between performance and training representation is otherwise inconsistent.

### COMRADE identifies novel hits for five targets

Following filtering of the ∼10.8 million-compound ZINC subset via CE-Screen and docking (Fig. 3a), 8-26 compounds were manually chosen for *in vitro* experimental testing against each target. This approach identified at least one previously untested ligand for each target, demonstrating inhibitory activities ranging from 4.10-124.60 μM, with a hit rate of 8-10% (Table 3). *In vitro* testing of EGFR, MEK1 and BRAF was carried out using immunoblotting, while M^pro^ activity was evaluated with a FRET-based enzymatic assay. The identified hits for EGFR and MEK1, while novel to these targets, have been previously reported to bind other proteins, highlighting their potential for repurposing. Notably, we discovered novel scaffolds for ligands targeting BRAF, M^pro^, and BCKDK, offering promising opportunities for further optimization and development.

**Figure 3:**
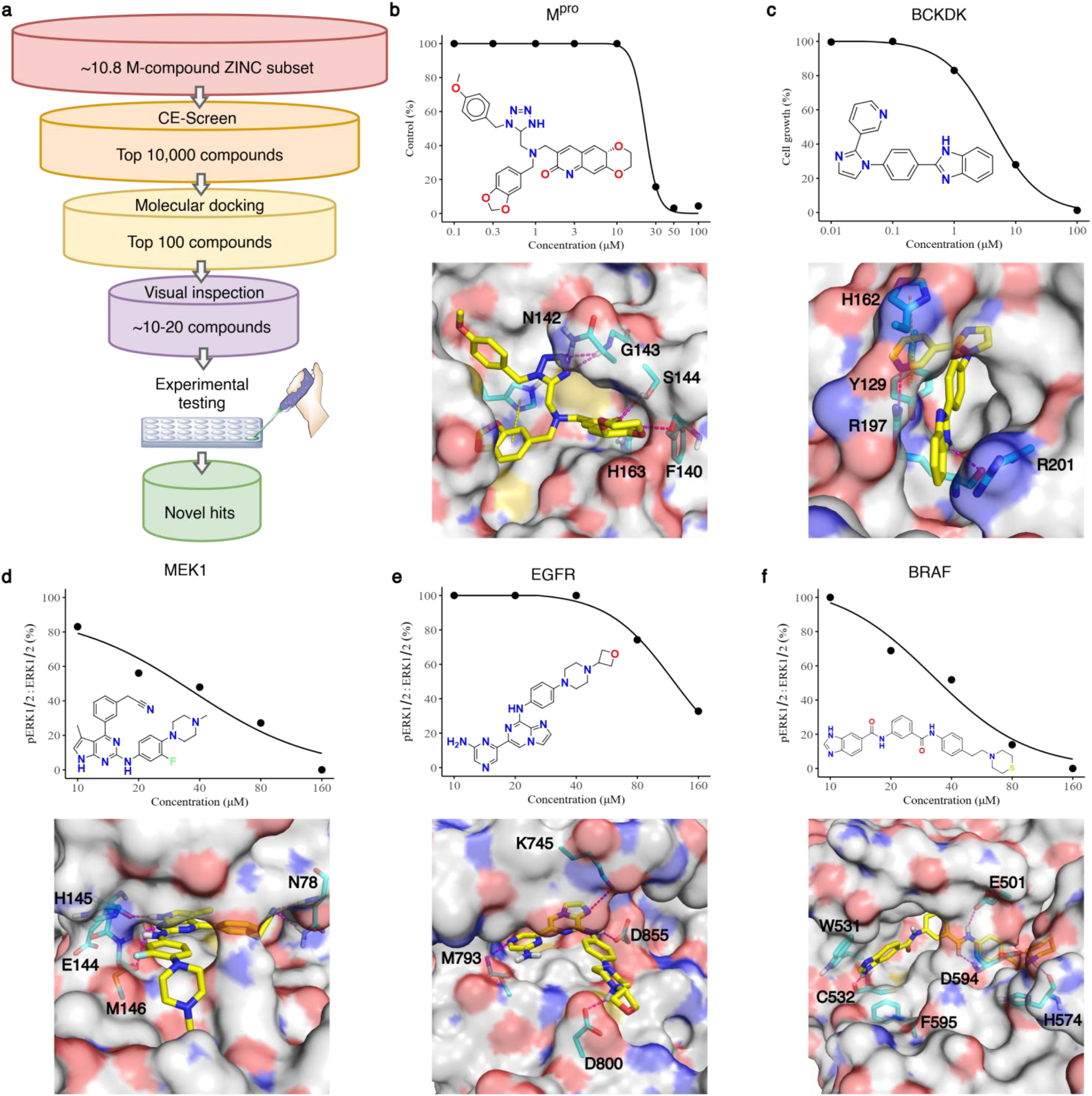
details of hits identified by COMRADE. **a**, the COMRADE framework workflow, as applied to screening ZINC for the five targets of interest. **b-f**, dose-response curves for the hits tested on their respective targets and the corresponding docking pose highlighting binding site interactions. **b**, FRET-based enzymatic assay of M^pro^’s hit against SARS-CoV-2 Mpro, with docking pose at the SARS-CoV-2 main protease binding site, showing hydrogen bonds with H163, the backbone of F140, and residues N142 and G143. **c**, Hep3B wildtype cells treated with BCKDK’s hit, with docking pose in the allosteric pocket of BCKDK, forming hydrogen bonds with R197 and engaging in π-π interactions with H162 and Y129, as well as cation-π interactions with R201. **d**, 293T transfectants expressing MEK1 treated with its hit, with docking pose at the ATP binding site of MEK1, displaying hydrogen bonds with E144, H145, M146, and N78. **e**, 293T transfectants expressing EGFR treated with its hit, with docking pose at the ATP binding site of EGFR, highlighting hydrogen bonds with K745, D855, M193, and D800. **f**, 293T transfectants expressing BRAF treated with its hit, with docking pose in the allosteric site of BRAF, featuring hydrogen bonds with C532, D594, E501, and H574, along with π-π stacking with W531 and F595, and cation-π interactions with H574.

Although BCKDK is now recognized as a high-value target in metabolic disease and oncology, the chemical space around it is still remarkably sparse: beyond the long-standing tool compound BT2^58^, only a handful of more potent chemotypes such as Pfizer’s thiophene PF-07208254^59^ and the repurposed antihypertensive valsartan have been described^60^, and each series is covered by active patent portfolios or suffers from mechanism-linked liabilities (e.g. mitochondrial uncoupling by BT2 or BCKDK-stabilization by some thiazoles). Recently, PF-07328948 has emerged as a promising BCKDK inhibitor and is current under Phase I clinical trials for HFpEF^65^, underscoring the translational potential of this target. Nonetheless, the limited diversity and accessibility of existing scaffolds present a significant bottleneck to broader therapeutic exploration. Driven largely by Pfizer and a few academic groups, current discovery efforts seek BCKDK inhibitors that can lower circulating branched-chain amino acids, improve insulin sensitivity and fatty-liver endpoints in obese or heart-failure models, and, more recently, suppress tumor growth and metastasis via MEK/ERK and focal-adhesion signaling^61–64^. Against this backdrop, our CE-Screen hit has generated an entirely new scaffold, and we set out to test the effects of this novel compound. We had previously generated several BCKDK KO Hep3B cell lines and profiled their expression levels of BCKDK (also known as BDK) via Western blot^66^. The novel compound was able to inhibit cellular proliferation in the BDK wild type cell line but not the BDK KO cell lines, indicating that the compound specifically targeted BCKDK and thereafter resulted in anti-proliferative effects (Fig. 4a). The novel compound demonstrated such effects consistently in other liver cancer cell lines (Fig. 4b-c) whilst showing negligible toxic effects in THLE-2 cell line which represents non-carcinogenic liver cells (Supplementary Fig. 3). All in all, its single-digit micromolar IC_50_ in Hep3B cells provides a compelling starting point for further medicinal chemistry optimization. Importantly, given the clear therapeutic rationale, active clinical development, and substantial industrial interest, the application of ML-driven approaches to explore new chemical space and accelerate hit identification for understudied yet high-value targets such as BCKDK is both timely and imperative.

**Figure 4:**
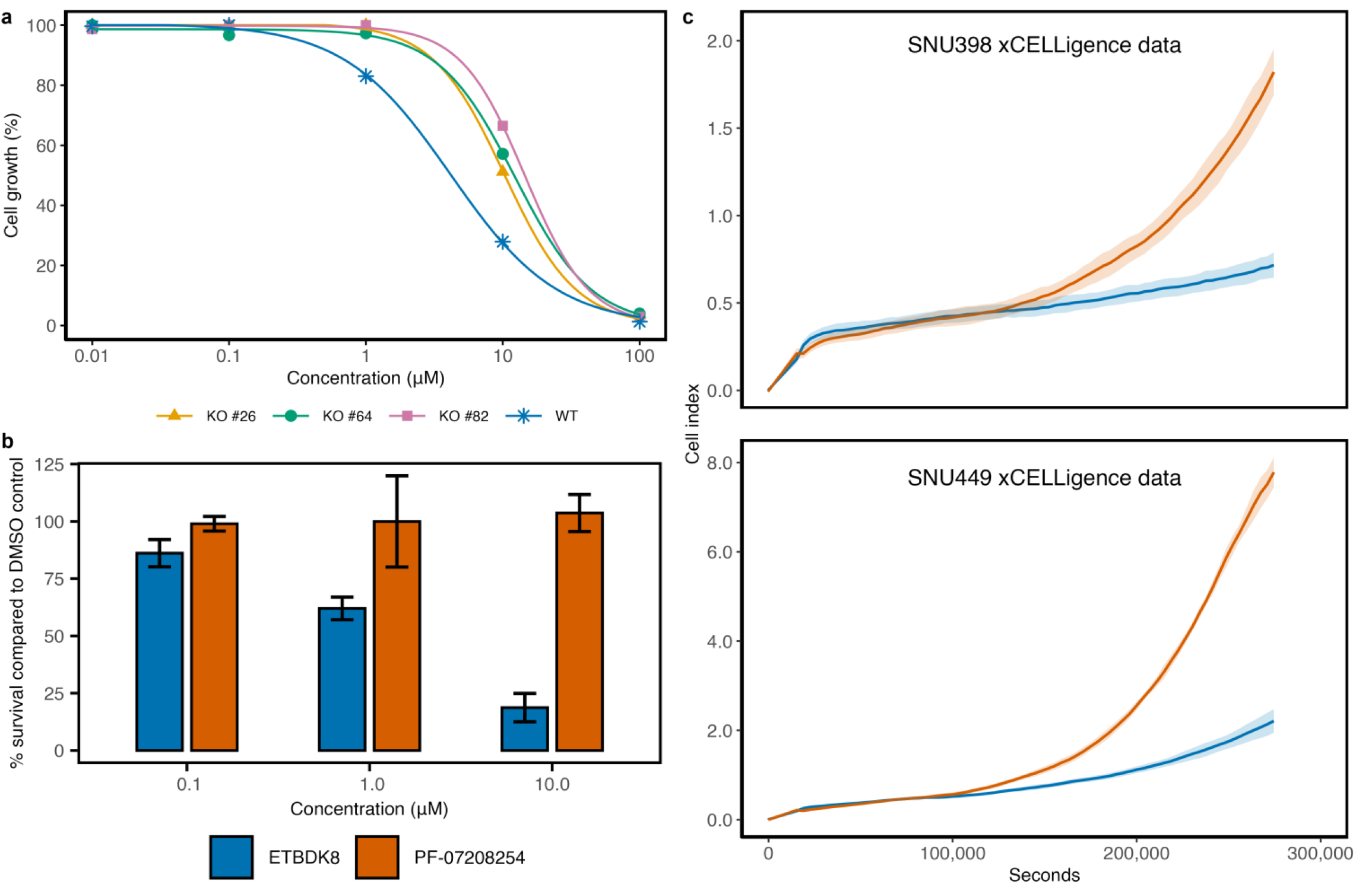
identification and characterization of the novel BCKDK inhibitor ETBDK8 in liver cancer cell models. **a**, validation of BCKDK dependence for ETBDK8 activity. Cell proliferation assays in BCKDK wildtype and knockdown Hep3B cell lines show that ETBDK8 displays BCKDK expression-dependent anti-proliferative activity, with reduced potency in knockdown cells. **b**, ETBDK8 outperforms the reference compound PF-07208254 in SNU182 cells across three tested concentrations. **c**, Real-time xCELLigence assays in SNU387 and SNU449 cells confirm that ETBDK8 shows superior potency over PF-07208254 at 10 µM, with reduced cell index progression relative to control.

### Each CE-Screen module is necessary and optimal

While CE-Screen consistently outperforms its competition, Table 4 considers whether its compression and classification modules are jointly necessary and optimal for its performance. This is done by replacing them with other modules and testing configurations where feature compression is disabled (Configurations 5-7), or replaced by a fully end-to-end architecture that couples compression directly with a neural network classification head (Configuration 4). Configurations employing gradient boosting^67^ classification heads are considered; gradient boosting serves as an alternate decision tree ensembling approach to Extra Trees and has recently been used with success for binding affinity prediction^68^. Also in comparison are configurations where TabNet replaces CE-Screen’s Extra Trees classification head. TabNet is a neural network specifically designed to address deep learning’s limitations with tabular data relative to decision tree-based methods through attentive feature selection^69^.

**Table 4:** logAUC performance of alternative CE-Screen configurations (best, <u>second best</u>). To assess variability in ML model training due to the random initialization of parameters and order of training data, testing was conducted four times with different random seeds. The mean and standard deviation of these runs are calculated and presented below.

| Configuration number | Feature compression | Classifier | BindingDB testing |  | ChEMBL testing |  |
| --- | --- | --- | --- | --- | --- | --- |
|  |  |  | Matched decoys | Random decoys | Matched decoys | Random decoys |
| 1 (CE-Screen) | Contrastive learning | Extra Trees | 49.69 ± 0.33 | **61.02 ± 0.14** | **55.44 ± 0.26** | **56.11 ± 0.11** |
| 2 | Contrastive learning | Gradient boosting | 47.04 ± 0.15 | 59.09 ± 0.35 | <u>54.42 ± 0.37</u> | 54.25 ± 0.75 |
| 3 | Contrastive learning | TabNet | 42.63 ± 0.95 | 56.38 ± 0.71 | 51.38 ± 0.43 | 52.29 ± 1.08 |
| 4 | End-to-end compression and classification neural network | | 42.54 ± 0.31 | 58.99 ± 0.07 | 53.47 ± 0.25 | <u>54.41 ± 0.11</u> |
| Classifier alone |  |  |  |  |  |  |
| 5 | None | Extra Trees | **52.97 ± 0.02** | 57.52 ± 0.02 | 49.53 ± 0.04 | 53.40 ± 0.03 |
| 6 | None | Gradient boosting | 49.86 ± 0.02 | <u>59.78 ± 0.01</u> | 51.19 ± 0.01 | 54.08 ± 0.03 |
| 7 | None | TabNet | 41.72 ± 0.83 | 57.94 ± 0.24 | 51.59 ± 0.62 | 52.07 ± 0.70 |
| Alternative feature compression |  |  |  |  |  |  |
| 8 | PCA | Extra Trees | <u>51.99 ± 0.05</u> | 57.34 ± 0.06 | 48.67 ± 0.10 | 51.17 ± 0.08 |
| 9 | PCA | Gradient boosting | 50.55 ± 0.00 | 56.86 ± 0.00 | 48.85 ± 0.00 | 52.35 ± 0.01 |
| 10 | PCA | TabNet | 39.27 ± 1.08 | 53.33 ± 1.36 | 46.95 ± 1.09 | 49.51 ± 2.55 |
| 11 | FA | Extra Trees | 51.98 ± 0.06 | 57.80 ± 0.02 | 48.91 ± 0.04 | 52.96 ± 0.05 |
| 12 | FA | Gradient boosting | 48.10 ± 0.03 | 58.71 ± 0.01 | 50.32 ± 0.00 | 53.32 ± 0.01 |
| 13 | FA | TabNet | 39.80 ± 2.37 | 54.85 ± 0.79 | 48.89 ± 0.64 | 49.70 ± 1.52 |

Across testing scenarios, CE-Screen achieves a 1.87-3.12% higher logAUC than the next-best configuration, with the exception of on BindingDB testing with matched decoys, where direct Extra Trees classification without compression yields a 6.60% higher logAUC. Notably, Configurations 8 and 11, which use Extra Trees classification with PCA- and feature agglomeration (FA)^70^-based feature compression (respectively) also outperform CE-Screen in this scenario by similar margins. This pattern suggests that CE-Screen may have slightly overfit to the D-DUD-E training data for this scenario relative to these simpler alternatives due to the additional complexity introduced by its use of neural network-based feature compression. However, in the other three scenarios, approaches employing neural network-based feature compression (Configurations 1-4) are approximately on par with or outperform these alternate Extra Trees configurations, with CE-Screen specifically achieving a 5.07-11.93% greater logAUC than they do. Overarchingly, therefore, CE-Screen’s inclusion of contrastive feature compression before classification improves performance. More specifically, that CE-Screen also consistently outperforms its end-to-end alternative by at least 3.12% across all scenarios – with a peak advantage of 16.81% on BindingDB with matched decoys – emphasizes the conceptual strength of CE-Screen’s particular use of contrastive feature compression explicitly decoupled from classification.

Nonetheless, it is still notable that most alternative CE-Screen configurations are also competitive with, and often outperform, the end-to-end models BridgeDPI, DrugBAN, PSICHIC, DLM-DTI, and MolTrans. This highlights the soundness of CE-Screen’s core design principle: to leverage rich, pre-computed biochemical embeddings from foundation models and domain-specific encodings rather than relying solely on task-specific end-to-end classification derived directly from simple compound-protein representations. Augmented with contrastive feature compression, CE-Screen shows itself to be a modular and scalable model that holistically outperforms state-of-the-art end-to-end pipelines, advancing the frontier of hybrid ML approaches for ligand discovery.

## Discussion

This work introduces COMRADE, a virtual screening framework that employs CE-Screen to rapidly triage large chemical libraries before targeted investigation with docking and experimental testing. COMRADE is successfully used to identify hits for targets that are both present in and significantly different from those encountered during its training. CE-Screen employs contrastive neural network-based feature compression to consolidate a diverse set of input representations, including protein language model and molecular embeddings, and a physics-based interaction potential encoding, into a compact, discriminative form optimized for use with its lightweight decision tree-based classifier. This hybrid, modular architecture departs from prevailing end-to-end deep learning approaches by decoupling representation learning from classification and incorporating expert-engineered features alongside learned embeddings.

CE-Screen consistently outperforms state-of-the-art end-to-end models on two challenging benchmarks, demonstrating greater robustness against overfitting and high ligand enrichment in imbalanced discovery scenarios. Beyond performance, CE-Screen’s modular design provides a forward-compatible platform: its feature compression can readily incorporate emerging biophysicochemical representations and alternative classifiers, enabling continuous improvement as ML and structural bioinformatics advance. More broadly, this decoupled compression-and-classification paradigm offers a architectural blueprint for accelerating other molecular prediction tasks – such as enzyme engineering, substrate selection, and antibody sequence optimization – to advance scalable discovery across bioinformatics and cheminformatics. Through its design and demonstrated generalization, COMRADE exemplifies a transferable approach to efficient, high-throughput compound-protein screening.

## Methods

### Retrospective training and testing datasets

#### Target and ligand compilation

Three datasets are employed in this work: a deduplicated version of the Directory of Useful Decoys, Enhanced^40^ (D-DUD-E), and original subsets of the BindingDB^41^ and ChEMBL^42^ databases. The datasets ligands (interacting compounds) are curated to prevent a small number of overrepresented, archetypal ligands from disproportionately influencing training or testing (Fig. 5a). Firstly, each ligand in the three datasets appears only once among them. In its original form, the Directory of Useful Decoys, Enhanced (DUD-E) dataset contains 1,998 ligands associated with 2-21 targets. This dataset is deduplicated into D-DUD-E by keeping only the single instances of these ligands associated with the target having the fewest overall number of ligands. Moreover, the ligand sets composing BindingDB and ChEMBL are made diverse (Fig. 5a): each ligand has an ECFP4 fingerprint^71^-based Tanimoto coefficient^72^ (T_c_) of no more than 0.70 and 0.50 (in BindingDB and ChEMBL, respectively) with any other ligand for the same target. T_c_ measures the structural similarity between two compounds as the ratio of their shared molecular substructure features to the total number of their unique features.

**Figure 5:**
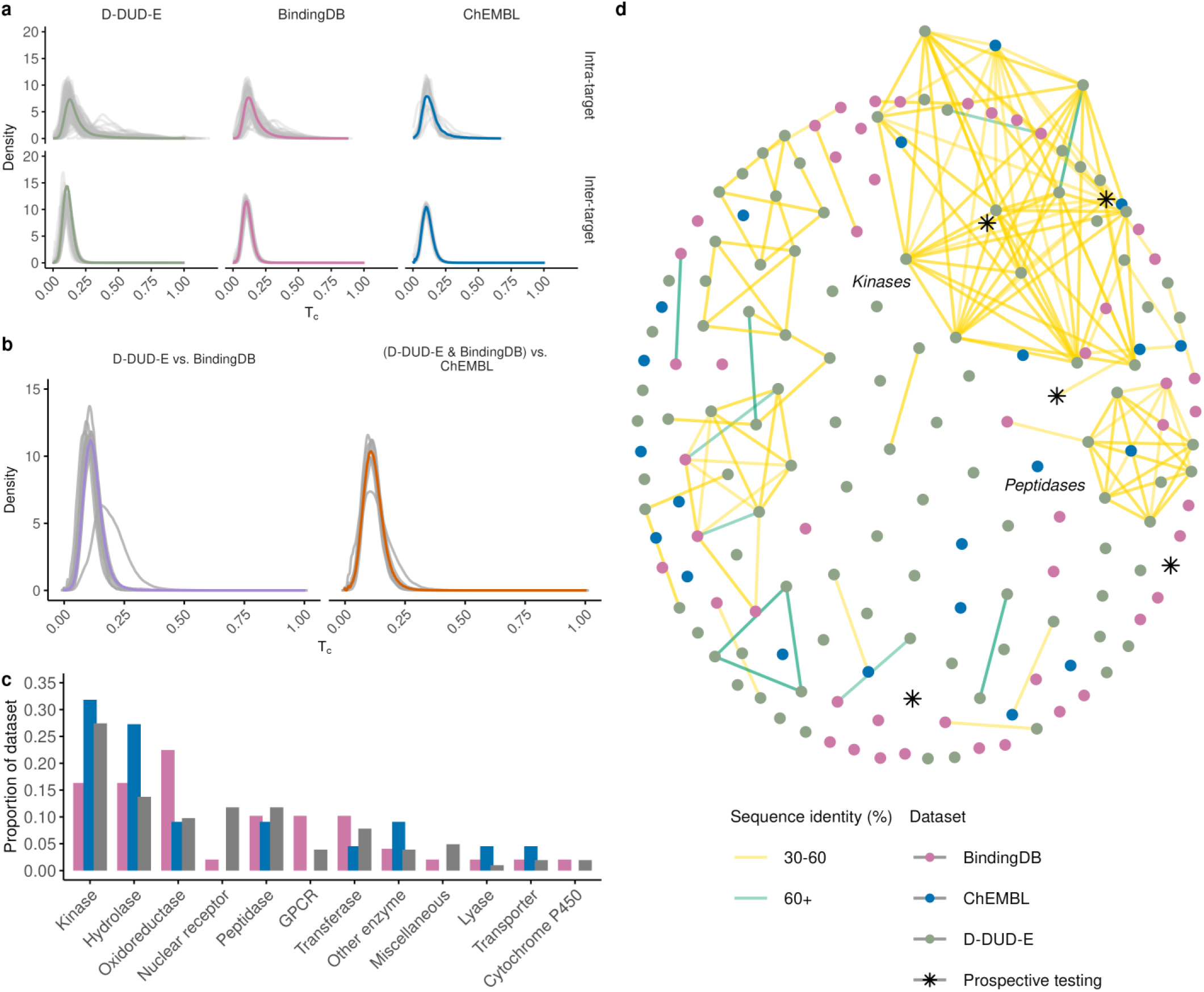
the targets and compounds that compose D-DUD-E, BindingDB, and ChEMBL. **a**, the datasets’ intra- and inter-target ligand ECFP4 fingerprint^71^-based Tanimoto coefficient^72^ (T_c_) distributions. T_c_ measures the structural similarity between two compounds as the ratio of their shared molecular substructure features to the total number of their unique features. Each dataset’s ligands, when compared with other ligands associated with the same target (intra-target), is 0.21 ± 0.06 (D-DUD-E), 0.18 ± 0.04 (BindingDB), and 0.13 ± 0.03 (ChEMBL) T_c_-similar. When these ligands are compared to ligands associated with other targets (inter-target), they are 0.11 ± 0.01 T_c_-similar in each dataset. **b**, the T_c_ distributions of ligands from the same target family compared between training and testing dataset pairs. For both pairs, D-DUD-E and BindingDB, and D-DUD-E and BindingDB (considered together) compared with ChEMBL, ligands from the same target family are 0.13 ± 0.02 T_c_-similar. **c**, the broad selection of protein families represented within the three datasets reflects their targets’ diverse biological roles and spans key functional groups, including signaling molecules, metabolic enzymes, regulatory proteins, and specialized catalytic families. **d**, the network of targets, represented as nodes. Edges are drawn between nodes if the identicality of the corresponding target pair’s sequences is greater than 30%, as calculated after performing pairwise global alignment between sequences using EMBOSS’s Needle program^74^. While most targets’ sequences differ significantly from others (being 8.75 ± 6.33% identical on average), about 50 targets, primarily from D-DUD-E and BindingDB, group into larger, mostly homogeneous clusters consisting of kinases or peptidases. These clusters are a consequence of their protein families’ highly conserved nature due to their respective structural and functional roles.

Ligands from BindingDB and ChEMBL exhibit a binding affinity of 1,000 nM or better (in terms of K_i_, IC_50_, K_d_, or EC_50_) with their target, with ChEMBL ligands specifically selected from assays that have the highest internal ChEMBL reliability score of 9^73^. Additionally, the targets that compose BindingDB and ChEMBL are chosen to represent the broad range of protein families also featured in D-DUD-E (Fig. 5c). Nonetheless, given that CE-Screen and other CPI models encode targets based on their sequences, targets are also selected to maintain relative sequence diversity within each dataset and distinctness between datasets (Fig. 5d, Supplementary Fig. 1). Moreover, the ligand sets associated with targets belonging to the same protein families (e.g., kinases) but from different datasets are also significantly distinct from each other in terms of the T_c_ (Fig. 5b, Supplementary Fig. 2).

#### Training dataset resampling

These target-ligand datasets are combined with decoy (non-interacting) compounds for their use in training and testing compound-protein interaction classification models. In their testing form, 50 decoys complement every ligand. However, when training on these datasets, decoys are under-sampled in a target-stratified random fashion to match the number of ligands. This parity ensures that ligands are not disregarded during model training. Without this adjustment, training on the original datasets could lead models to focus disproportionately on target-decoy pairs, as their numerical superiority would dominate the optimization of model parameters aimed at minimizing total training loss.

#### Random decoy selection

Random decoys are selected from the ZINC database^43^ to serve as one of the two kinds of non-interacting examples of compound-protein pairs, complementing the interacting pairs compiled from the chemical databases used in this work (Fig. 6). Each target’s decoy set is constructed tranche by tranche, drawing from the ZINC tranches comprising compounds positive logP values. This procedure is iterative: compounds are randomly sampled from the tranche being cycled to until one compound with an T_c_ of at most 0.30 with any previously selected decoy compound is found. Then, the other tranches are each similarly sampled until a decoy set with 50 times the number of interacting compounds (ligands) is constructed. Since this construction is performed independently for each target, any decoy compounds inadvertently duplicated across decoy sets are removed and randomly replaced by unique compounds from the 90 tranches.

**Figure 6:**
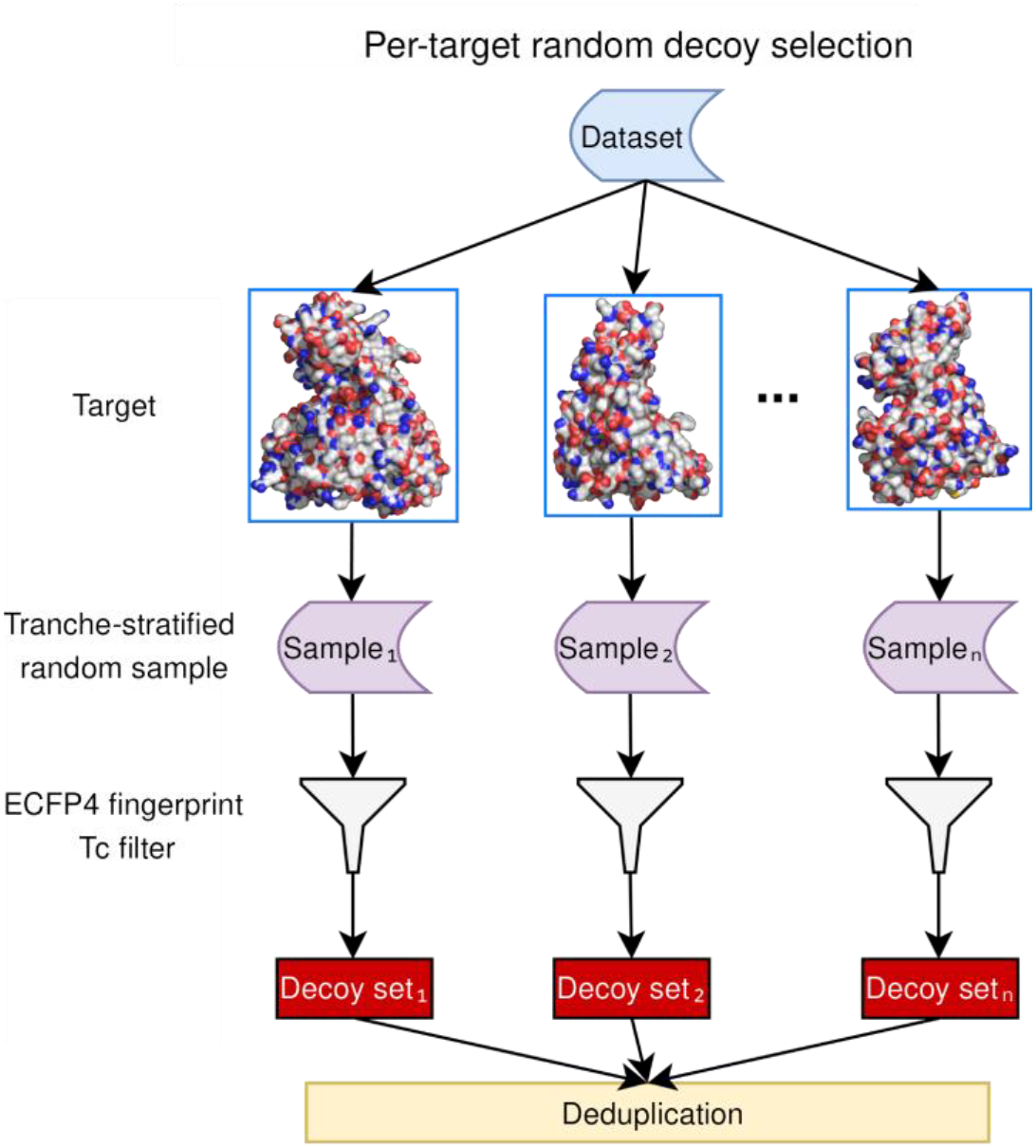
the random decoy selection procedure. Decoys are assigned to targets from a drug-like subset of the ZINC database in a physicochemically- and structurally-diverse manner. Decoys duplicated across targets are replaced by unique, randomly-selected ZINC compounds.

### The COMRADE framework

#### Creation of target representations

CE-Screen uses two target representations created by the DLM-LSTM^24^ and ESM-2^21^ protein Language Models (pLMs). These pLMs process a target’s amino acid sequence with *r* residues to create embedding matrices **T**_DLM−LSTM_ ∈ ℝ^*r*×6165^ and **T**_ESM−2_ ∈ ℝ^*r*×1024^ respectively. Each row of these matrices represents the embedding of its corresponding amino acid. Both matrices are aggregated into embedding vectors **t**_DLM−LSTM_ ∈ ℝ^6165^ and **t**_ESM−2_ ∈ ℝ^1024^ respectively by taking a weighted column-wise mean. The weight *w*_*k*_ ∈ (0, 2] associated with the values of row *k* ∈ {1, …, *r*} is set according to the minimum distance *d*_*k*_ ∈ ℝ^+^ Å from the corresponding amino acid residue to the target’s co-crystal ligand (Fig. 7). This distance is found with reference to the target’s 3-dimensional Protein Data Bank^75^ (PDB) resolved structure.

**Figure 7:**
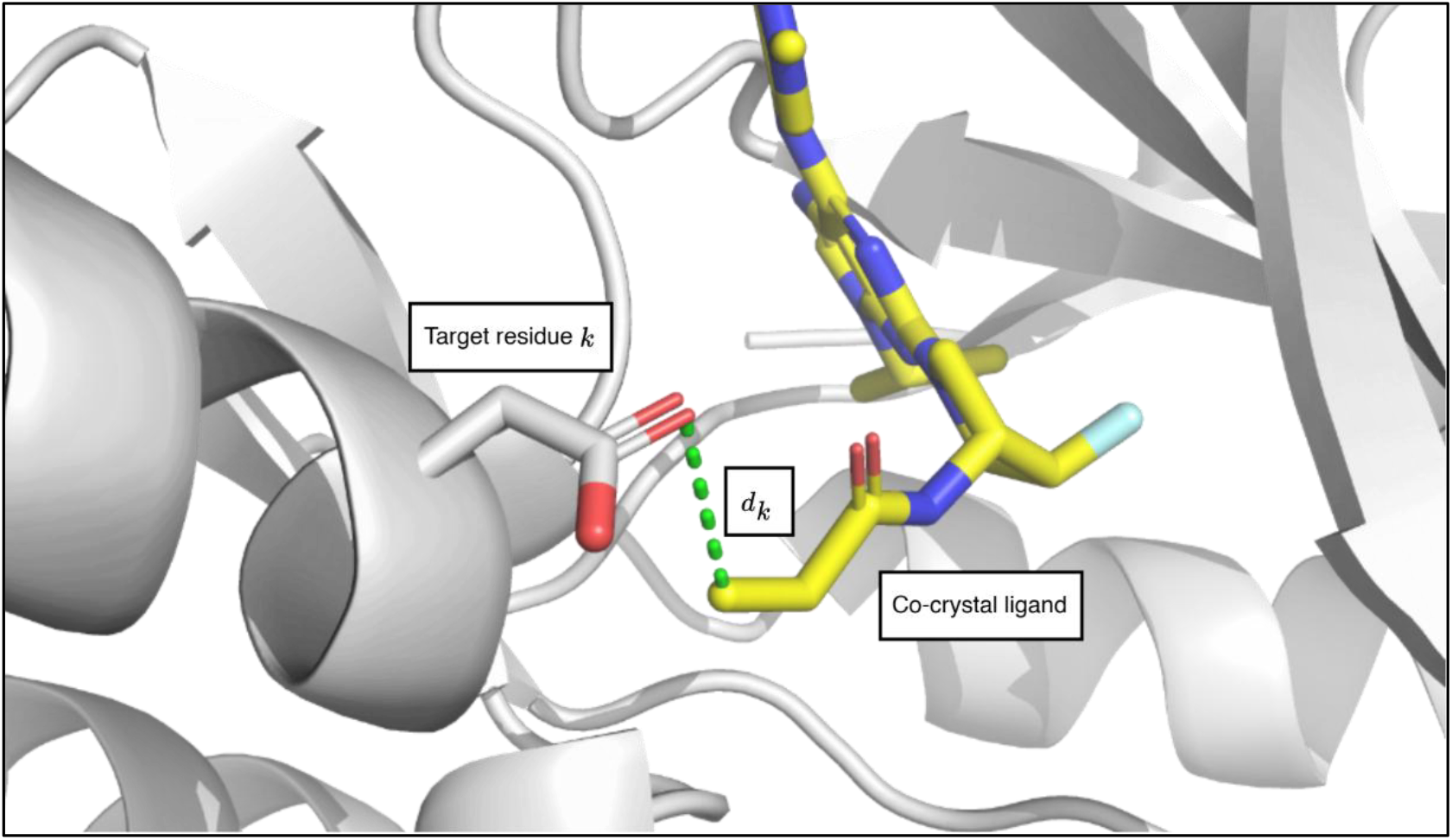
the minimum distance *d*_*k*_ between residue *k* and the target’s co-crystal ligand. The protein embedding matrices **T**_DLM−LSTM_ and **T**_ESM−2_ are aggregated into embedding vectors **t**_DLM−LSTM_ and **t**_ESM−2_ respectively via a weighted column-wise mean. The weight assigned to each row *k* (representing amino acid residue *k*) is set according to *d*_*k*_, the minimum distance between the residue and the target’s co-crystal ligand within its 3-dimensional PDB resolved structure. In the displayed example, *d*_*k*_ = 3.67 Å is the length of the dashed line between the helical residue’s delta oxygen atom and the ligand’s carbon atom, and represents the shortest straight-line distance between the residue and ligand.

The weight is then set as follows:

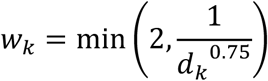

The specific form of this function, along with other hyperparameters relevant to the creation of CE-Screen’s input representation, cannot be directly optimized using the training dataset through a learning algorithm. Instead, they are adjusted by tuning based on a hold-out validation dataset derived from the training dataset. This empirical process involves exploring a predefined set of hyperparameter value ranges, evaluating their impact on model performance, and selecting the combination that yields the best results on the validation set. These hyperparameters are further detailed in Supplementary Table 1.

After matrix aggregation, **t**_DLM−LSTM_ is compressed into 50 dimensions using feature agglomeration^70^ via the `scikit-learn`1.3.2 Python package^76^. Feature agglomeration uses hierarchical clustering to identify groups of features that behave similarly across targets’ embeddings, and aggregates feature values in each group by taking their mean.

#### Creation of compound representations

CE-Screen uses 4 compound representations created based on their SMILES string form^23^. Mordred descriptors^25^ and mol2vec embeddings^26^ are calculated using the `deepchem`2.8.0 Python package. MACAW embeddings^27^ are calculated using the `macaw_py`1.0.1 Python package with the parameters `n_components`(controlling the embedding dimensionality) and `n_landmarks`(controlling the number of compounds used as landmarks by the method) set to 15 and 100 respectively. Lastly, MolFormer-XL embeddings^24^ are calculated using the `ibm/MolFormer-XL-both-10pct`model from the `transformers`4.45.2 Python package^77^.

#### Creation of interaction potential encoding (IPE)

An IPE is created in four steps for each compound-protein pair on the basis of their atoms’ Merck Molecular Force Field (MMFF) types (of the 94s variant^29^). First, MMFF types are assigned to each heavy atom in the target’s binding pocket and in the compound using the `RDKit`2024.03.5 Python package^78^, with the atoms comprising the pocket defined as those 10 Å or closer to the target’s co-crystal PDB ligand. Next, the MMFF types of the atoms covalently bonded to these atoms are also identified. This step enables the creation of an MMFF type pair for each atom in the binding pocket and in the compound, where the first element of the pair represents the atom’s own type and the second represents the types of its neighboring atoms. In the third step, the frequency of unique combinations of target and compound MMFF type pairs is calculated. Lastly, pair counts are consolidated into a 300-dimensional vector via principal component analysis using the `scikit-learn`1.3.2 Python package.

### Deep contrastive feature compression

#### Contrastive learning network architecture

The target, compound, and IPE representations are concatenated into **h**_0_, a 2,457-dimensional real-valued vector whose dimensions are median-centered and rescaled according to their interquartile ranges using the `RobustScaler`functionality within the `scikit-learn`1.3.2 Python package. This overall representation is compressed into 750 dimensions using a residual neural network^33^ employing contrastive representation learning (Fig. 1b). This network consists of two sequential components: a feature-processing module and an embedding module. As with the hyperparameters relevant to the creation of CE-Screen’s input representation, those concerning this network’s modules and CE-Screen’s classification head are also empirically tuned via validation and are listed in Supplementary Table 2.

The first module transforms **h**_0_ into a higher-dimensional hidden representation **h**_normalized_ ∈ ℝ^2650^, and the second compresses **h**_normalized_ into a parsimonious form **v**_2_ ∈ ℝ^750^ designed for interaction classification. The contrastive learning network is implemented using the `torch`2.4.1 Python package^79^.

Specifically, the feature-processing module comprises of two feedforward layers, each consisting of a linear transformation followed by a Sigmoid Linear Unit (SiLU) activation function^80^, defined as:

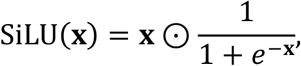

where **x** ∈ ℝ^*k*^, *k* ∈ ℕ. To enhance gradient flow and improve model convergence, residual skip connections are also incorporated. These connections add the layer’s input to its output. The computation at each layer *i* ∈ {1, 2} can therefore be expressed as:

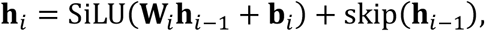

where **h**_*i*_ ∈ ℝ^2650^ is the hidden representation created by layer *i* ; 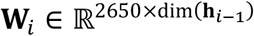 and **b**_*i*_ ∈ ℝ^2650^ are respectively the weights and biases of the linear transformation at layer *i*; skip(**h**_i−1_) ∈ ℝ^2650^ is the output of the skip connection, defined as:

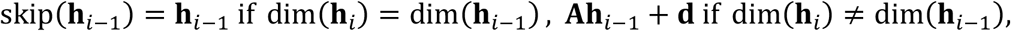

where **A** ∈ ℝ^2650×2457^ and **d** ∈ ℝ^2650^ transform the layer’s input into a form that can be added to its output. This transformation is required only for layer 1, where the input and output dimensions (2,457 and 2,650 respectively) are mismatched. In contrast, both the output of layer 1 (i.e., the input to layer 2) and the output of layer 2 are 2,650-dimensional. Before being passed on to the embedding module, the hidden representation **h**_2_ ∈ ℝ^2650^ undergoes ℓ_2_-normalization along the feature dimension:

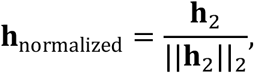

constraining the processed features to have unit norm by projecting them onto the surface of a hypersphere. This normalization ensures that the hidden representations have a consistent scale, which improves numerical stability during training and emphasizes angular relationships, better allowing for the clustering of same-class representations through supervised contrastive representation learning.

The embedding module compresses **h**_normalized_ ∈ ℝ^2650^ into a 750-dimensional representation designed for compound-protein interaction prediction by CE-Screen’s classification head. It achieves this by using two feedforward layers with linear transformation, SiLU activation, and skip connections. Unlike the feature-processing module, however, the first of this module’s two layers produces a 750-dimensional real-valued hidden representation **v**_1_, and the second layer further processes it into the network’s output, **v**_2_ ∈ ℝ^750^. The computation at each layer *j* ∈ {1, 2} can therefore be expressed as:

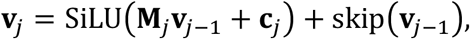

where **v**_0_ := **h**_normalized_; **v**_*j*_ ∈ ℝ^750^ is the representation created by layer *j*; 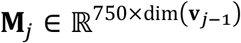 and **c**_*j*_ ∈ ℝ^750^ are respectively the weights and biases of the linear transformation at layer *j*; skip(**v**_*j*−1_) ∈ ℝ^750^ is the output of the skip connection, defined as:

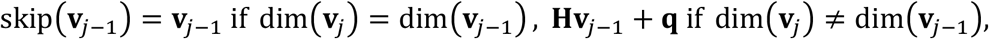

where **H** ∈ ℝ^750×2650^ and **q** ∈ ℝ^750^ transform the layer’s input into a form that can be added to its output. This transformation is required only for layer 1, where the input and output dimensions (2,650 and 750 respectively) are mismatched. In contrast, both the output of layer 1 (i.e., the input to layer 2) and the output of layer 2 are 750-dimensional.

#### Contrastive learning network optimization

The weights and biases associated with both the linear transformations and the skip connections are learnable parameters, optimized during training to minimize a modified supervised contrastive loss function calculated over training data. This optimization enables the network to adapt to the training dataset and improve the quality of the learned representations for interaction prediction using CE-Screen’s classification head. Specifically, the training objective encourages the contrastive learning network to structure its output embedding space such that ligands’ and decoys’ embeddings are geometrically distinct.

The objective is minimized through mini-batch gradient descent, a method that updates the learnable parameters by computing gradients over small subsets of the training dataset. While this approach is computationally efficient compared to those that do so over the entire dataset or individual examples^81^, the stochasticity inherent in sampling mini-batches can introduce noise into the gradient estimates, potentially leading to suboptimal convergence and poorly generalizing embeddings. A heterogeneous batching strategy is therefore employed to ensure each mini-batch contains ligands and decoys associated with a diverse set of targets. This approach consistently encourages the network in its training to learn to create holistic ligand-decoy embedding separations that generalize well across targets.

Specifically, let the training dataset of size *N* be represented as:

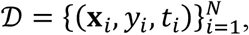

where **x**_***i***_ ∈ ℝ^2457^ denotes the *i* -th example’s input feature vector; *y*_*i*_ ∈ {0, 1} is its interaction class label (0: decoy, 1: ligand); *t*_*i*_ is the corresponding target’s PDB identifier (the dataset contains *M* ≤ *N* unique targets). To ensure target diversity within each mini-batch, 𝒟 is shuffled such that examples from different targets alternate as uniformly as possible. This shuffling process interleaves examples associated with distinct targets, creating a new ordering that avoids consecutive sequences of examples from the same target. Mini-batches of size *B* (in this work, 512) are then constructed sequentially from the shuffled dataset, with each mini-batch therefore containing examples from a diverse set of targets.

The ⌈*N*/*B*⌉ mini-batches constituting the dataset are trained over 350 times (i.e., in 350 epochs). A modified form of the supervised contrastive loss function is calculated and backpropagated through the network after it is applied to each mini-batch. Specifically, the modified loss function used in this work combines a supervised contrastive loss term^34^ with an additional decorrelation regularization term. This modification encourages the embeddings created by the trained contrastive learning network to not only be discriminative between ligands and decoys, but also less redundant for better use with CE-Screen’s Extra Trees classification head.

Namely, the supervised contrastive loss term guides the network to cluster examples within the same interaction class (ligands or decoys) together within in the embedding space, and to drive examples from different classes apart. Let **f**_*i*_ ∈ ℝ^750^ denote the embedding of the *i* -th example. The supervised contrastive loss term is then defined as:

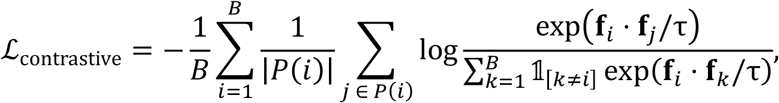

where *P*(*i*) = {*j* | *y*_*i*_ = *y*_*j*_ *for j* ≠ *i*} is the set of examples from the mini-batch within the same interaction class as *i*; 𝟙 is the indicator function; τ is the temperature parameter (in this work, set to 0.60), which scales similarity scores to control the sharpness of the loss probability distribution; and the dot product quantifies embedding similarity. The decorrelation regularization term penalizes redundancy in the embedding space by encouraging the minimization off-diagonal entries in the embeddings’ covariance matrix. Let the matrix of embeddings in the mini-batch be denoted as **F** ∈ ℝ^*B*×750^. The decorrelation term is then given by:

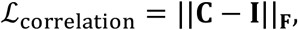

where:

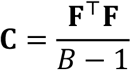

is the covariance matrix of the features, **I** is the identity matrix, and || · ||_*F*_ denotes the Frobenius norm. The total loss for a mini-batch is then computed as:

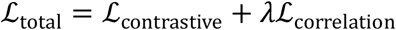

where *λ* is a hyperparameter controlling the weight of the decorrelation regularization. In this work, *λ* is set to 1.

The learnable parameters of the network are updated using the Adam optimizer^82^ with a learning rate of 5 × 10^−7^. They are initialized using a simplified Xavier normal initialization^83^. Namely, for a linear layer with *n*_input_ input features, the weights are drawn from a normal distribution:

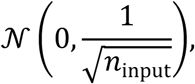

and the biases are initialized to 0. This initialization ensures that the initial magnitude of the gradients remains stable across layers during backpropagation, facilitating efficient convergence.

### Classification head

#### Classification head architecture and optimization

CE-Screen uses an Extra Trees^30^ classification head as implemented by the `scikit-learn`1.3.2 Python package. This head consists of 2,500 individual decision trees, each of which independently classifies the input compound-protein pair as either interacting (output: 1) or non-interacting (output: 0). The final classification is determined through a majority voting mechanism across the outputs of all the trees. The mean of the constituent decision trees’ votes also provides a pseudoprobability of interaction, reflecting the proportion of trees predicting an interaction. This score is not a calibrated probability but is used within the COMRADE framework to prioritize compounds for further evaluation. Only pairs with the highest scores are selected for downstream investigation and interpretation via docking using the Glide-SP program in its standard precision mode^37^.

Decision trees are hierarchical constructs that partition their input space into regions corresponding to different output classes. These partitions are formed by recursively splitting decision nodes, where each split divides the data based on a feature and threshold that optimize a specified criterion. The minimum number of compound-protein examples required to perform a split at a node is set to 2. The quality of each potential split is determined using the Gini impurity criterion, which evaluates the weighted average of the Gini impurities of the two resulting child nodes. Let *n*_left_ and *n*_right_ denote the number of examples in the left and right child nodes, respectively, and *n*_total_ = *n*_left_ + *n*_right_ denote the total number of examples in the parent node. The Gini impurity for the split is given by:

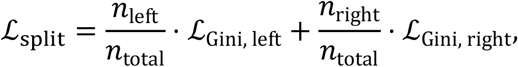

where the Gini impurity for a single node is:

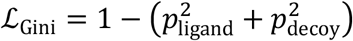

and *p*_ligand_ and *p*_decoy_ represent the proportions of examples belonging to each class within the node. The Gini impurity ranges from 0 (pure split) to 0.5 (completely mixed split). At each node, the Gini impurity is evaluated for all potential splits, and the split minimizing ℒ_split_ is selected.

During training, the Extra Trees classification head optimizes the ensemble as a whole by randomly sampling subsets of the training dataset and input feature dimensions to construct each tree. This sampling ensures high variance among individual trees, allowing the ensemble to reduce overfitting by leveraging several independent decision-making processes into a robust final prediction.

### Prospective testing

#### Molecular docking

The protein structure of five protein targets the serine/threonine-protein kinase B-Raf (BRAF, PDB ID: 1UWH), epidermal growth factor receptor (EGFR, PDB ID: 2RGP), and Mitogen-activated protein kinase (MEK1: 1S9J), SARS-CoV-2 main protease (M^pro^, PDB ID: 7QBB), and the branched-chain ketoacid dehydrogenase kinase (BCKDK, PDB ID: 8EGQ) were obtained from the PDB database. The protein structure was prepared by adding hydrogen atoms using Protein Preparation Wizard^84^. CE-Screen was used to assign a score to the each of the ∼10.8 million ZINC compounds. The top-ranked 10,000 compounds for each target were prepared and optimized using the LigPrep module^36^, which generated all possible ionization states at pH 7.0 ± 2.0 and multiple conformations for each compound. A receptor grid box was generated around the bound ligand using the Receptor Grid Generation tool in Glide^37–39^. The grid box dimensions were set to encompass the entire active site with a sufficient margin for accurate docking. The receptor grid was designed to allow flexibility in ligand positioning during the molecular docking simulations. The prepared compound database was docked into the binding site of target protein using the Glide Standard Precision (SP) docking method. For kinases, BRAF, EGFR, MEK1 and BCKDK, docked to both the ATP and allosteric binding sites. The docking results were analyzed, and compounds were ranked based on their Glide-SP Score values, reflecting the predicted binding affinities. The top 100 compounds were visually inspected and shortlisted the compounds for further experimental investigation.

#### M^pro^ enzymatic assay

The SARS-CoV-2 M^pro^ fluorescence resonance energy transfer (FRET) substrate Dabcyl-KTSAVLQ/SGFRKME (Edans) was synthesized, and the M^pro^ enzymatic assays were carried out exactly as previously described in pH 6.5 reaction buffer containing 20 mM HEPES (pH 6.5), 120 mM NaCl, 0.4 mM EDTA, 20% glycerol, and 4 mM DTT^85,86^. The assay was performed in 96-well plates with 100 μL of 100 nM M^pro^ protein in the reaction buffer. Then 1 μL testing compound at various concentrations was added to each well and incubated at 30 °C for 30 min. The enzymatic reaction was initiated by adding 1 μL of 1 mM FRET substrate (the final substrate concentration is 10 μM). The reaction was monitored in a Cytation 5 image reader with filters for excitation at 360/40 nm and emission at 460/40 nm at 30 °C for 1 h. The initial velocity of the enzymatic reaction with and without testing compounds was calculated by linear regression for the first 15 min of the kinetic progress curve. The IC_50_ was determined by plotting the initial velocity against various concentrations of the compounds using the following equation in Prism 8 software:

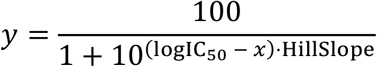

where *x* is the logarithm of inhibitor concentration, and *y* is the normalized enzyme velocity.

#### BRAF, MEK1 and EGFR activity assays

Plasmids that encode BRAF, constitutively active MEK1 mutant (DD), or EGFR were transfected into 293T cells by using lipofectamine 2000 (Invitrogen). 24 hours later, 293T transfectants were replated into 6-well plates, and then treated with testing compounds for two hours. Afterwards, 293T cells were harvested and lyzed with RIPA buffer supplemented with protease inhibitors, kinase inhibitors and phosphatase inhibitors to prepare whole cell lysates as in our previous studies^87,88^. The phospho-ERK1/2 as well as BRAF, MEK1(DD) or EGFR in whole cell lysates were analyzed by SDS-PAGE together with immunoblottings. Antibodies used in these assays include anti–phospho-ERK1/2 (no. 4370, Cell Signaling Technology); anti-FLAG (F3165) and anti-β-actin (A2228) (Sigma-Aldrich); anti-ERK1/2 (A0229, AB clonal); and horseradish peroxidase-labeled secondary antibodies (the Jackson laboratory). All antibodies were diluted following the manufacturers’ recommended protocols.

#### BCKDK assay

To validate the effectiveness of these compounds in-vitro experiments were conducted. We have performed cell viability assays in both Hep3B wildtype (ATCC) and BCKDK knockdown cell lines to assess if these compounds are specific and potent. The Hep3B BCKDK knockdown cell lines have been previously generated with the methods reported in our previous publication^89^. The cells were maintained in Dulbecco’s modified Eagle’s medium (Gibco). All media were supplemented with 10% fœtal bovine serum (Gibco) and 1% penicillin-streptomycin (Gibco). All cells were cultured in a humidified incubator at 37°C and 5% CO_2_.

For the cellular viability assays, we have employed sulforhodamine B (SRB) assay. Briefly, the cells were seeded into 96-well tissue-culture plates at a density of 1000 cells per well. At the end point of the experiment, the medium is aspirated and replaced with 100 µL cold 10% trichloroacetic acid (TCA) into each well and thereafter, incubation at 4 °C for 1 h takes place. The plates were then washed with distilled water and dried in 37 °C for 30 min, before 80 µL of SRB dye (Sigma Aldrich) was added to each well for 5 min. The plates were then washed with 1% acetic acid, dried and thereafter, the dye was then resolubilized in 10 mM Tris. The plates were then recorded for the absorbance at a wavelength of 510 nm, with the values expressed in arbitrary units. All SRB assay experiments were performed in triplicates.

To test the phosphorylation levels of BCKDHA upon 24 hours of respective compound treatment, Hep3B cells were lysed in buffer containing 20 mM Tris-HCl (pH 7.5), 150 mM NaCl, 2 mM EDTA, 1.5 mM MgCl2, 1% NP-40, protease inhibitor cocktail, phosphatase inhibitors (NaF, NaVO3 and PMSF). After the lysed samples were rotated at 4°C for 1 h and centrifuged at 16000g for 15 minutes, supernatant was obtained and quantified using Protein BCA assay (ThermoFisher). 4X LDS with β-mercaptoethanol buffer was then added to samples and heated at 95°C for 5 minutes before subjecting to electrophoresis using Bio-Rad’s MiniPROTEAN system. After separation, protein lanes were transferred onto a nitrocellulose membrane using wet transfer at a constant 300mA for 110 minutes. Membrane was subsequently washed with 0.1% TBS-Triton (TBST) before standard blocking procedure. The antibodies used are as follows: Anti-BCKDHA (phospho S293) antibody (Abcam, #ab200577), GAPDH Antibody (6C5) (Santa Cruz Biotechnology, #sc-32233) and BCKDK (Cell Signaling, #82318).

## Supplementary Information

**Supplementary Table 1:** compound-protein representation-related hyperparameters. The 2,457-dimensional representation input into CE-Screen consists of individual representations encoding the target, the compound, and the compound-protein interaction encoding. six hyperparameters, primarily controlling the dimensionality of these individual representations, are tuned through validation on a held-out subset of the training dataset.

| Related to | Hyperparameter | Considered values | Selected value | Remarks |
| --- | --- | --- | --- | --- |
| Target representations | Constant compared to in $w_k$ formulation | {1, 2, 3, 4} | 2 | Controls the maximum weightage a residue’s embedding has on the target’s overall sequence embedding. A greater value allows for a greater weight. |
| | Exponent of $d_k$ in $w_k$ formulation | {0.25, 0.50, 0.75, 1.00, 1.25} | 0.75 | Residues farther away from the co-crystal ligand have lower weight than hyperparameter (1.). A greater value of this exponent makes this this weight reduce more quickly with $d_k$. |
| | Feature agglomeration dimensionality of $\mathbf{t}_{\text{DTM-LSTM}}$ | {25, 50, 75, 100, 200, 300, 400, 500} | 50 | Controls DTM-LSTM embeddings’ output dimensionality; a greater value increases the influence of the representation on CE-Screen’s output, and therefore performance. |
| Compound representations | Number of MACAW components | {15, 30, 45, 60, 100, 200, 300, 400} | 15 | Controls the MACAW embeddings’ output dimensionality. |
|  | Number of MACAW landmarks | {10, 50, 100, 200, 300, 400, 500, 1000, 2000} | 50 | Landmark compounds are selected from the dataset to act as fixed points, with their pairwise distances determining how all other compounds are positioned in the output embedding space. |
| Interaction potential encoding | Number of interaction potential principal components | {50, 100, 200, 300, 400, 500, 600, 700, 800} | 300 | Controls encodings’ output dimensionality. |

**Supplementary Table 2:** CE-Screen’s hyperparameters. CE-Screen is comprised of two parts: a contrastive learning network that compresses a compound-protein pair’s input representation into a 750-dimensional embedding, and an Extra Trees classification head that uses this embedding to create an interaction prediction. Both parts have parameters and decision rules (respectively) that they dynamically learn from the training dataset. In contrast, their hyperparameters are not directly optimized through the training process but are preset. The values of these hyperparameters are tuned through validation on a held-out subset of the training dataset.

| CE-Screen module | Hyperparameter | Considered values | Selected value | Remarks |
| --- | --- | --- | --- | --- |
| Contrastive learning network | RobustScaling input representation | {No, Yes} | Yes | Controls whether the network’s inputs are median-centered and IQR-scaled. |
|  | Batch size | {128, 256, 512, 768, 1024} | 512 | The number of examples processed together in a single forward and backward pass of training. |
| | Learning rate | $\{1 \times 10^{-7}, 5 \times 10^{-7}, 1 \times 10^{-6}, 5 \times 10^{-6}, 1 \times 10^{-5}, 5 \times 10^{-5}\}$ | $5 \times 10^{-7}$ | Controls the size of the adjustments made to the model’s weights during each backward pass. |
| | Temperature ($\tau$) | {0.10, 0.25, 0.40, 0.50, 0.60, 0.90, 1.00} | 0.5 | Controls the scaling of the dot product (similarity measure) between embeddings within the supervised contrastive loss function, with lower temperatures making calculated loss more sensitive to embedding dissimilarity. |
|  | Parameter initialization | {Xavier Uniform, Xavier Normal, Simplified Xavier, He, Kaiming Normal} | Simplified Xavier | The method that sets the initial values of the network’s weights and biases. |
| | $\mathcal{L}_{\text{correlation}}$ multiplier ($\lambda$) | {0.10, 0.50, 1.00, 1.50, 2.00} | 1 | Controls the influence of output embeddings’ correlatedness (relative to their supervised contrastive discriminativeness) in network training. |
|  | Number of feature-processing hidden layers | {1, 2, 3, 4} | 2 | The number of hidden layers used to process feature representations before embedding. |
|  | Number of embedding hidden layers | {1, 2, 3, 4} | 2 | The number of hidden layers used to refine the embedding itself. |
|  | Activation function | {Rectified Linear Unit (ReLU), Leaky ReLU, SiLU} | SiLU | The non-linear function applied to layer outputs, affecting representational capacity and gradient flow. |
|  | Use skip connections | {No, Yes} | Yes | Controls whether to include shortcut connections between layers to mitigate vanishing gradient issues and accelerate learning. |
| | $\ell_2$-normalization | {No, Yes} | Yes | Controls whether the output of the feature-processing module of the network is normalized before being passed on to the embedding module. |
|  | Embedding dimensionality | {250, 500, 750, 1000} | 750 | The dimensionality of the embedding module’s output. |
| Extra Trees classification head | Number of trees | {500, 1000, 1500, 2000, 2500, 3000} | 2500 | The number of independent decision trees comprising the classification head. |
|  | Minimum number of samples required to split an internal node | {2, 4, 6, 8, 10} | 2 | The number of samples required to split a decision tree’s internal node (that is, further partition the tree’s input space). |
|  | Split criterion | {Gini, Entropy, Log loss} | Gini | The metric used to evaluate the quality of internal node splits. |

## Supplementary Figures

**Supplementary Figure 1:**
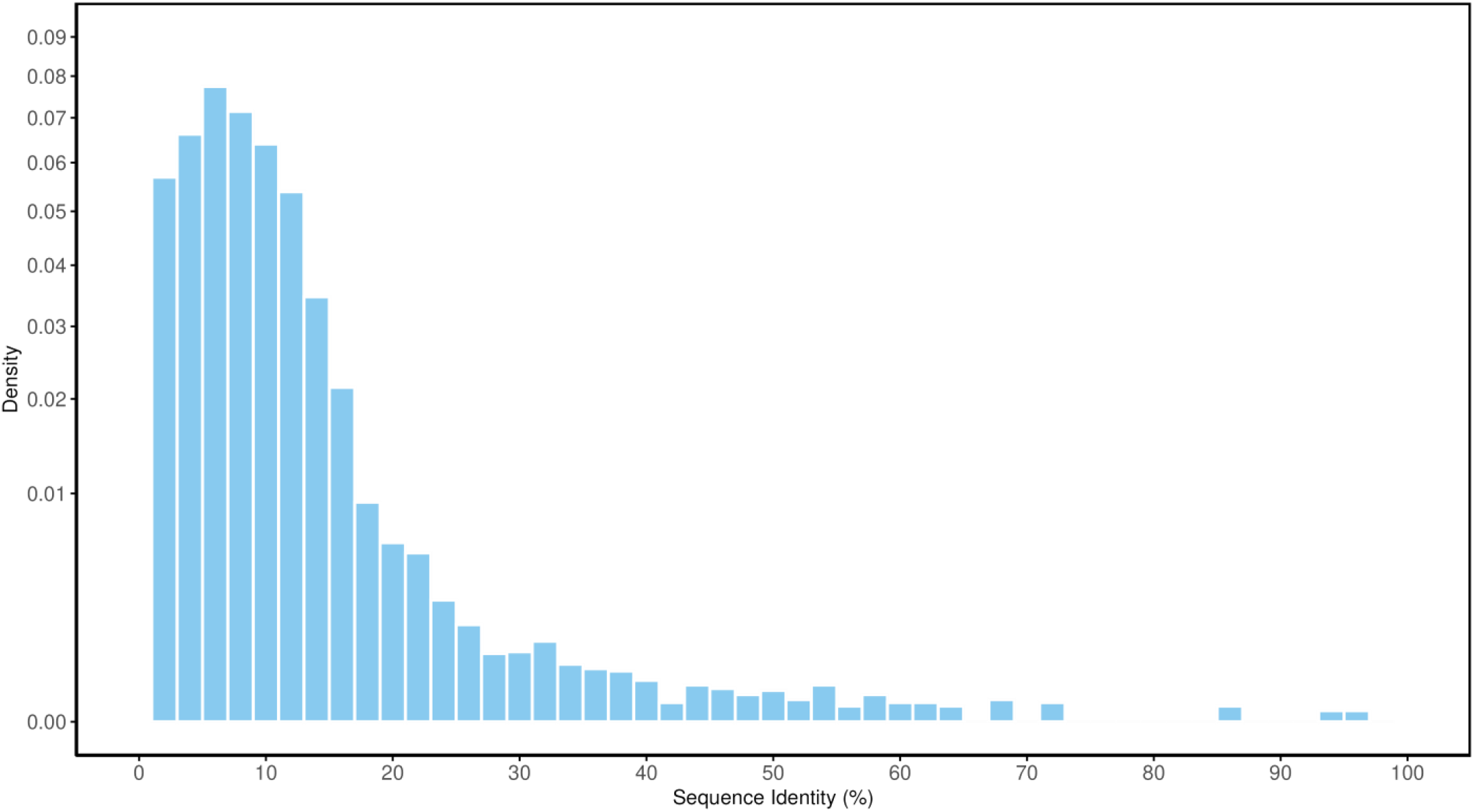
the distribution of target sequence identicality. Target sequences from all three datasets, after pairwise global alignment, are on average 8.75 ± 6.33% identical. Targets more than 20% identical to each other usually belong within clusters of targets with conserved sequences such as kinases, nuclear receptors, peptidases, or GPCRs.

**Supplementary Figure 2:**
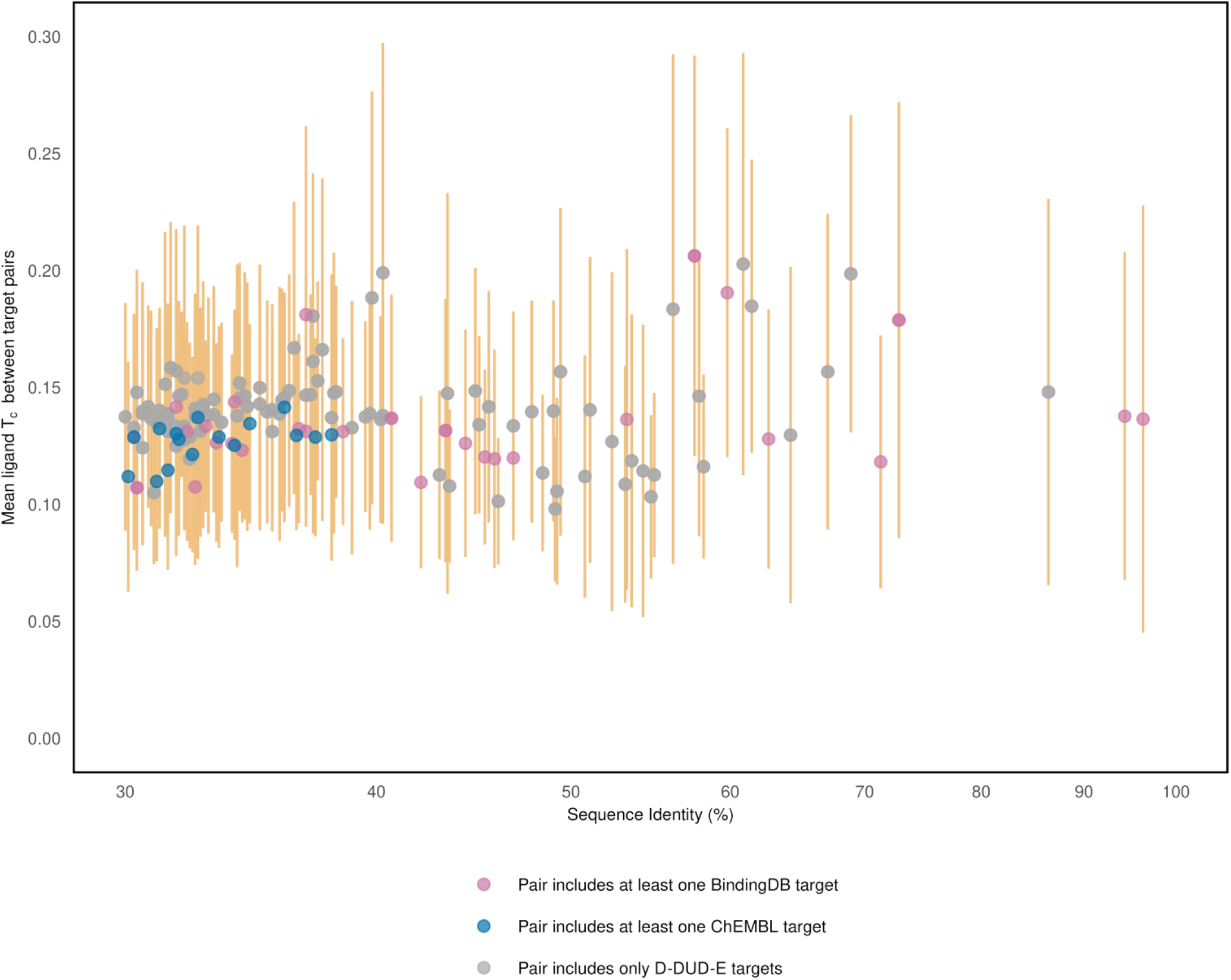
the relationship between mean ligand T_c_ and target identicality for target pairs with greater than 30% sequence identity. The mean of the pairwise T_c_ values of ligands associated with closely-related targets is low, ranging from 0.10-0.21 (the T_c_ values’ standard deviation is represented by orange bars.

**Supplementary Figure 3:**
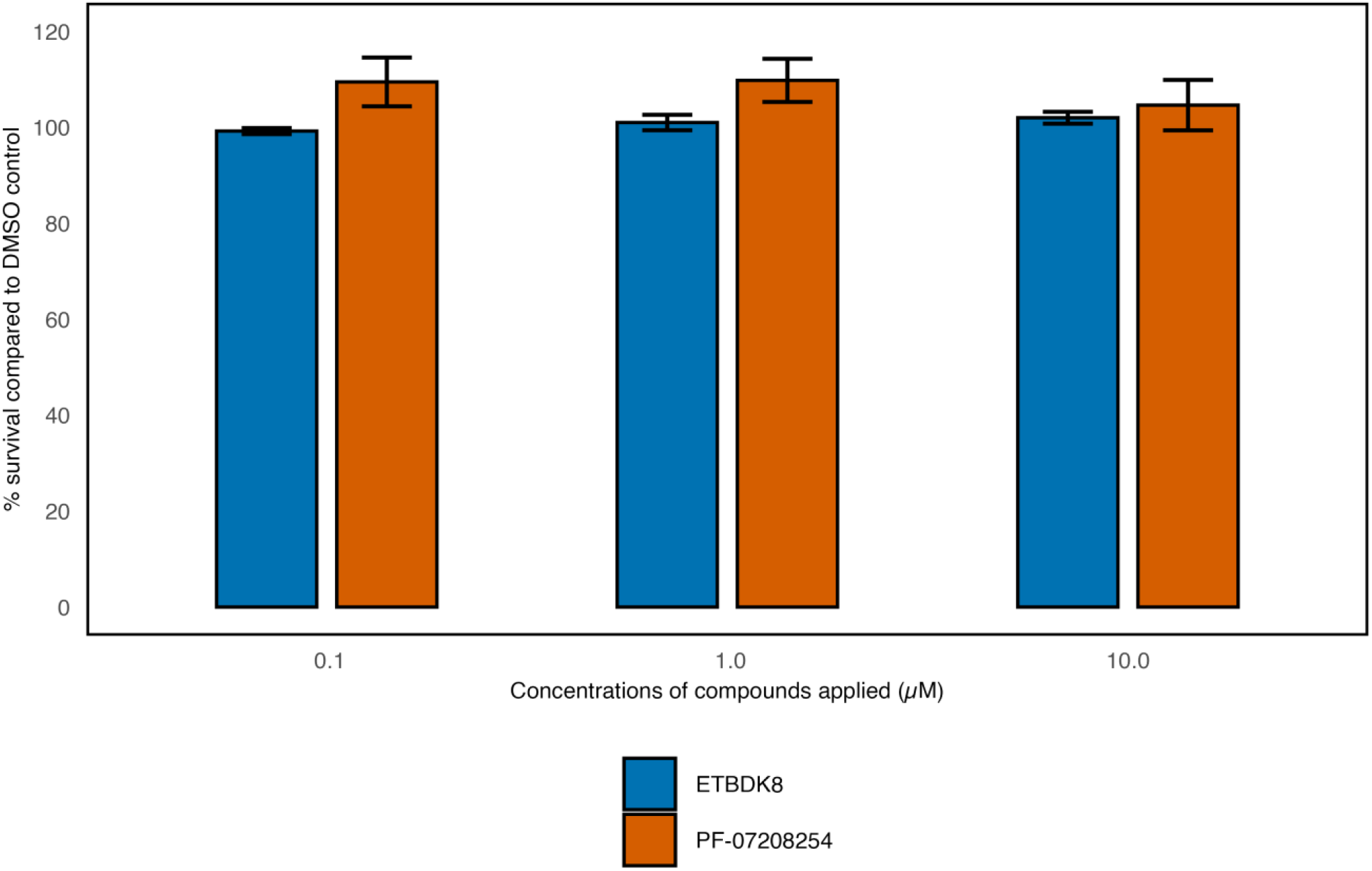
ETBDK8 shows minimal toxicity in THLE-2 cell line. ETBDK8 does not cause significant toxic effects in the THLE-2 cell line, which is an epithelial liver cell line commonly used for comparison with liver cancer cell lines.

